# Biophysical simulations of fMRI responses using realistic microvascular models: insights into distinct hemodynamics in humans and mice

**DOI:** 10.1101/2025.10.08.680976

**Authors:** Grant Hartung, Avery J. L. Berman, Sava Sakadžić, Andreas Linninger, David A. Boas, Jonathan R. Polimeni

**Affiliations:** Athinoula A. Martinos Center for Biomedical Imaging, Massachusetts General Hospital, Charlestown, MA, USA; Department of Radiology, Harvard Medical School, Boston, MA, USA; Department of Physics, Carleton University, Ottawa, ON, Canada; Institute of Mental Health Research, Royal Ottawa Mental Health Centre, Ottawa, ON, Canada; Department of Bioengineering, University of Illinois at Chicago, Chicago, IL, USA; Department of Neurosurgery, University of Illinois at Chicago, Chicago, IL, USA; Neurophotonics Center, Department of Biomedical Engineering, Boston University, Boston, MA, USA; Harvard-MIT Program in Health Sciences and Technology, Massachusetts Institute of Technology, Cambridge, MA, USA

**Author notes:** Corresponding author: Jonathan R. Polimeni, Richard M. Lucas Center for Imaging Department of Radiology, Stanford University,1201 Welch Road, Stanford, CA 94305, USA. Richard M. Lucas Center for Imaging, Department of Radiology, Stanford University, Stanford, CA, USA.

**Keywords:** high-resolution fMRI, neurovascular coupling, functional neuroimaging, cerebrovasculature, vascular architecture, microvascular anatomy, fMRI physics, computational modeling, vascular synthesis, finite element analysis, computational fluid dynamics, CMRO_2_, functional hyperemia

## Abstract

Functional Magnetic Resonance Imaging (fMRI) is broadly used to measure human brain activity, however the hemodynamic changes that comprise the fMRI response to neuronal activity are often interpreted using microscopy data in mice. These microscopy data provide ground-truth observations of how individual blood vessels respond to neuronal activity and thus form the basis of our fundamental understanding of neurovascular coupling. Although these invasive experiments provide invaluable insight, there are striking differences in the vascular architecture of mouse and human brains that may influence the hemodynamic response. Motivated by this, we developed a biophysical modeling framework for realistic hemodynamic simulations in both mouse and human cerebral cortex. For this, we utilized Vascular Anatomical Network (VAN) models that explicitly represent the full microvascular tree as a single connected network, originally based on anatomical reconstructions from a given location of mouse cerebral cortex. We extended the VAN modeling framework using synthetic VAN models representing the microvascular network at a single location of the human cerebral cortex. To account for larger size and complexity of the human VAN models, we developed an efficient computational framework to simulate the full hemodynamic responses in this human model and compared the simulated fMRI responses between mice and humans. Our biophysical simulations are based entirely on first principles (e.g., conservation of mass); model parameter values were fixed across all simulations, not tuned to fit data, as they represent meaningful physical constants taken from previous measurements. Only two simple calibrations were tuned for each simulation, to match baseline perfusion rates (blood flow) and oxygen extraction (OEF). Our results show that differences in microvasculature indeed influenced the hemodynamic response and led to observable differences in timing—e.g., the simulated fMRI response peak in humans was delayed by ∼2 s compared to that of mice, consistent with prior fMRI observations. While there are many known differences in vascular architecture in rodents and humans, we also discovered that, unexpectedly, an asymmetry in the numbers of branches of the penetrating intracortical arterioles and venules appears to be conserved across species. We demonstrate through further simulations that this anatomical property may also be needed for suitable hemodynamic responses. Our framework thus provides a valuable tool for bridging *in-vivo* microscopy of microvascular dynamics to human fMRI.

## 2 Introduction

Functional Magnetic Resonance Imaging (fMRI) is widely used to non-invasively measure neuronal activity and has contributed substantially to our understanding of the human brain. Of the functional neuroimaging methods available for routine use in humans, fMRI can uniquely measure function dynamically across the entire brain. The most common fMRI signal is based on the blood oxygenation level–dependent (BOLD) contrast due to its robustness and sensitivity. This contrast, however, reflects a complex interplay between several aspects of the hemodynamic response that accompany brain activity. BOLD responses represent changes in blood flow, volume, oxygenation and neural metabolism that all change with neuronal activity, providing an indirect measurement of brain function. Recent studies, however, have demonstrated that hemodynamics within the smallest blood vessels are more tightly coupled to neuronal activity than previously believed (Devor et al. 2003; Nizar et al. 2013; Chen et al. 2011; Cho et al. 2022; Poplawsky et al. 2015; Boido et al. 2019). This suggests better localization (in space *and* time) of underlying neural signals—and more information regarding the underlying neuronal activity of interest—could potentially be inferred from human fMRI with a more complete understanding of the relationship between neuronal and vascular dynamics. Currently, this relationship, termed neurovascular coupling, is primarily studied using *in-vivo* microscopy in small animal models, which provides direct, ground-truth observations of how individual blood vessels respond to neuronal activity. While these measurements are often motivated by a desire to improve the interpretation of human fMRI data, what is missing is a means to link the observable BOLD fMRI signals measured in humans back to microscopic hemodynamic changes observable only in these small animal models.

Realistic biophysical simulations could potentially help bridge between scales as well as bridge between imaging modalities to relate these fine-scale hemodynamic changes to the measured BOLD signal (Polimeni and Lewis 2021; Griffeth and Buxton 2011; Gagnon et al. 2015; Mandeville et al. 1999; Scheffler et al. 2021; Mester et al. 2024; Epp et al. 2020; Báez-Yáñez et al. 2017). Such simulations would provide a platform for understanding how microvascular anatomy and dynamics together shape macroscopic hemodynamics and the BOLD fMRI signal. This approach would also allow for testing of assumptions (implicit and explicit) made about the complex vascular response to neuronal activity and the underlying interplay of blood flow, volume, and oxygenation changes. Thus, this framework can be used to test our understanding of the hemodynamic response and identify knowledge gaps. These models further allow *in silico* experimentation that is not possible *in vivo*, such as investigating how variations of individual anatomical or physiological parameters affect the resulting BOLD response, providing deeper insights into how specific aspects of the vascular anatomy and physiology influence fMRI measurements. We note, however, that this framework is most powerful when it is used in tandem with empirical studies—i.e., when the model inputs and validations are taken directly from experimental data, and when the model can generate testable hypotheses and guide experimental design.

Previous biophysical models articulated our understanding of how changes in blood flow, volume and oxygenation give rise to the BOLD fMRI response at coarse spatial scales. (Polimeni and Lewis 2021; Griffeth and Buxton 2011; Gagnon et al. 2015; Mandeville et al. 1999; Scheffler et al. 2021; Báez-Yáñez et al. 2017). Classic BOLD simulation approaches, such as those using the Balloon Model framework (Buxton et al. 2004; Mandeville et al. 1999; Davis et al. 1998), transform *macroscopic* changes in blood flow and oxygen metabolism into *macroscopic* BOLD responses at the spatial scale of an fMRI voxel. This framework accurately captures BOLD dynamics across impressively wide ranges of experimental conditions (Buxton 2012). Using these models, distinct temporal features—such as the initial dip or post-stimulus undershoot—can be linked to transient decoupling between blood flow, blood volume, and oxygen consumption (Griffeth and Buxton 2011). This modeling framework, however, was originally derived from the low-resolution hemodynamic data available at the time. Often, the inputs to these models, such as estimates of blood flow or perfusion changes accompanying neuronal activation, are taken from fMRI data (e.g., Arterial Spin Labeling), which are themselves difficult to interpret. This is one key limitation of such models, since a lack of confidence in the inputs leads to even less confidence in the outputs. Moreover, the insights gained from many simulations is further hindered by the lack of realism of the underlying microvascular anatomy and physiology (Buxton 2012). Furthermore, the Balloon Model framework does not lend itself to modeling non-BOLD fMRI contrasts (Huber et al. 2019) which are increasingly used yet are in some cases may be difficult to interpret in terms of the underlying vascular physiology and/or hemodynamics.

Biophysical models based on simplified representations of individual blood vessels, such as random-cylinder models (Boxerman et al. 1995; Bandettini and Wong 1995; Uludağ et al. 2009; Ogawa et al. 1993), have also provided an understanding of how vascular geometry impact the fMRI signal. However, the lack of connectivity between vessels in these random-cylinder models precludes investigation of hemodynamics through the vascular network.

These limitations grow in importance as the imaging resolution of human fMRI improves and prior assumptions are revisited, and recent Balloon Model extensions have begun to address how vascular architecture influences hemodynamics. In high-resolution fMRI, adjacent voxels can sample from different levels of the vascular hierarchy and the hemodynamics of neighboring voxels are coupled across cerebral cortical depths (Havlicek and Uludağ 2020; Markuerkiaga et al. 2016). Increasing detail of the interdependence of hemodynamics—such as the hemodynamics within capillaries and downstream veins—offers new insights, such as how neuronal activity within a specific cerebral cortical layer manifests as observed patterns of BOLD responses across cortical depths. Modeling approaches with even more realistic vascular anatomy and interdependencies of hemodynamics have simulated observations of suppressed hemodynamic responses in the region immediately surrounding the site of activation (Boas et al. 2008; Huppert et al. 2007; Devor et al. 2003; Lorthois et al. 2011). These studies demonstrated how meaningful aspects of BOLD responses can be captured only when blood flow and oxygenation are coupled through a connected vascular network. These models, however, still use abstract simplifications of real vascular geometry which has a more complicated, interconnected structure (Blinder et al. 2013; Gould et al. 2017; Schmid, Tsai, et al. 2017). To fully benefit from ground-truth measures of microvascular dynamics and relate these to the measured fMRI signal will require more realistic descriptions of microvascular anatomy and physiology.

To integrate more realistic vascular interconnectivity and geometry, a new class of biophysical models has recently been introduced termed “Vascular Anatomical Network” (VAN) modeling (Boas et al. 2008; Fang et al. 2008; Gagnon et al. 2015) that explicitly represents all blood vessels at a single cortical location using reconstructions from optical microscopy data (Gagnon et al. 2015; Schmid, Tsai, et al. 2017; Linninger et al. 2013; Hartung, Badr, Moeini, et al. 2021; 2021; Genois et al. 2021; Pfannmoeller et al. 2020; 2021; Linninger et al. 2019; Payne and Lucas 2018; Park and Payne 2016; Gould et al. 2017; Gould and Linninger 2015; Báez-Yáñez et al. 2025; Báez-Yáñez and Petridou 2024; Lorthois et al. 2011). VAN modeling represents hemodynamics by taking direct measurements of diameter changes of individual vessels responding to neuronal activity as inputs—made possible by modern *in-vivo* microscopy technology. It then combines fluid dynamics, oxygen transport kinetics and magnetic field changes to compute the resulting BOLD fMRI signal, offering enhanced physiological interpretability and concreteness relative to previous models.

VAN models derived for mouse cortex have been used to predict an inter-relationship between baseline blood velocity, hematocrit, and cortical depth that was later validated *in vivo* (Hartung et al. 2018; Cheng et al. 2019). VAN models extended to simulating hemodynamic responses to neuronal activity predicted unexpected BOLD response characteristics that were later confirmed with conventional- and high-resolution human fMRI (Gagnon et al. 2015; Viessmann et al. 2019). For example, the BOLD response amplitude was found to vary with the local angle between the surface normal of the cortex and the main magnetic field of the MRI scanner. This bias could only be discovered using models with explicit representations of realistic vascular geometry. This raises the question of whether VAN models explicitly representing human brain vasculature can better predict features of human fMRI data.

While rodents are a common experimental model for understanding fMRI signals in humans, their vascular *topology* (connectivity and hierarchy) differs substantially from humans. For instance, there is an approximate 1:3 ratio of intracortical descending arterioles to intracortical ascending venules in mouse but a 2.1:1 ratio in humans (Schmid, Barrett, et al. 2017). A different balance between arteries and veins could potentially influence hemodynamics and the BOLD response in meaningful ways.

Vascular *geometry* also differs between species, including vessel densities, diameters, lengths, and the overall size of the interconnected microvascular hierarchy (Hartung, Badr, Mihelic, et al. 2021; Cassot et al. 2006; Lauwers et al. 2008; Cassot et al. 2009; Blinder et al. 2013; Schmid, Barrett, et al. 2017). Cortical thickness in humans is ∼2–4 times larger than in mice, translating to longer distances from the pial surface to the capillary bed. The extent to which differences in angioarchitecture result in differences in BOLD response timings (Lambers et al. 2020) remains largely unknown. This creates a need to understand the limits of using rodent vascular anatomical data to model angioarchitecture in humans, and whether microvascular hemodynamics differ in humans as a result.

Human VAN simulations could address this gap in knowledge and answer many open questions, however there are substantial technical challenges to achieve this. The current mouse *anatomical* input data were derived from invasive imaging methods (Blinder et al. 2013; Fang et al. 2008; Gagnon et al. 2015) not suitable for humans. While valuable reconstructions of human brain vasculature have been derived from histological data using ink injections (Cassot et al. 2006), the volumes are too thin to capture the vascular topology needed to model blood flow patterns. Additionally, these reconstructions are only available from small cortical regions where ink injection succeeded. Modern three-dimensional microscopic imaging based on tissue clearing may be capable of extending the imaging volume in human brain specimens (Chung and Deisseroth 2013; Bernier et al. 2019), however uniformly staining and reconstructing the full microvascular network accurately remains challenging. Furthermore, measurements of individual microvessels responding to neuronal activity, used as the required *physiological* input data, are also unavailable in humans. There are also computational challenges to simulating a human-sized VAN, which would be >125-fold larger than previous mouse VAN models used for BOLD simulations (Gagnon et al. 2015; Genois et al. 2021).

Here, we present an updated computational framework for biophysical simulations of the BOLD response for both mice and humans. To address the missing anatomical data needed for human modeling, we extended our existing vascular network synthesis method (Linninger et al. 2019; Hartung, Badr, Mihelic, et al. 2021) to generate an anatomically accurate human VAN. To address the missing physiological data, we extrapolated existing rodent data (Tian et al. 2010; Uhlirova et al. 2016) to humans. For computational tractability, we extended our scalable oxygen transport model (Hartung, Badr, Moeini, et al. 2021) that can handle the larger human VAN size and complexity. Our biophysical simulations are based entirely on first principles (e.g., conservation of mass); the model parameter values are fixed across all simulations, not tuned to fit data, as they represent meaningful physical constants taken from previous measurements. Only two simple calibrations were tuned for each simulation, to match baseline perfusion rates (blood flow) and oxygen extraction (OEF). Using this framework, we successfully simulated human BOLD response dynamics and tested our hypothesis that differences in vascular architecture between humans and mice lead to known differences in BOLD response dynamics. We found that differences in vascular geometry between humans and mice, related to differences in cortical thickness, may contribute to the slower BOLD response in humans. Unexpectedly, we also observed that differences in vascular topology may be associated with reduced passive venous ballooning in humans during activation. Finally, we discovered that the observed asymmetric branching of intracortical arterioles and venules may be required for a realistic BOLD response amplitude and that this topological feature of the microvasculature appears to be shared by mice and humans. Given the complexity of our framework, and the challenges associated with implementation, we also provide the source code, source data, simulation data, and a convenient graphic user interface to aid in widespread use of these methods.

## 3 Methods

Simulation of a human VAN required three methodological advancements. The first is a modified VAN synthesis algorithm used to generate the human VAN anatomy. The second is an updated computational platform to handle the large human VAN model. The third is extrapolation of arterial dilation measurements from the mouse cortex to the thicker human cortex. We then compare the BOLD responses between mouse and human.

### 3.1 Synthesizing VAN models

A key hurdle to simulating a human VAN is the lack of suitable microvascular microscopy data for a sufficient VAN reconstruction. To overcome this, we employ an image-based Cerebrovascular Network Synthesis (iCNS) algorithm (Linninger et al. 2019; Hartung, Badr, Mihelic, et al. 2021) based on Constrained Constructive Optimization (Karch et al. 2000) to generate synthetic VANs (sVANs) that closely mimic real reconstructed VANs (rVANs) that are directly derived from microscopy data, as summarized in **Fig. 1**. We selected this method for its ability to create VANs that are amenable to hemodynamic simulations from statistical distributions of common topological or geometric properties (Linninger et al. 2019; Hartung, Badr, Mihelic, et al. 2021). This method first creates an initial artery and vein before adding new segments until the desired density is reached. Segments are added stochastically to the existing trees following strict geometric constraints creating a bifurcation at an optimized position (Linninger et al. 2019; Hartung, Badr, Mihelic, et al. 2021). The different stages of vessel generation orient the vessels using different criteria—starting with new vessels aligned tangential to the cortical surface (e.g., pial vessels), then aligned radially (e.g., intracortical penetrating vessels), then oriented randomly (e.g., capillaries)—to capture the distinct microvascular hierarchical levels, as shown schematically in **Fig. 1B**. This generates a fully connected vascular network that is compatible with our dynamic blood flow simulations, described below. The final stage overwrites the initial geometric properties of each segment (i.e., segment diameter and tortuosity) such that the distribution of final geometric properties matches those taken from real microvascular data (Hartung, Badr, Mihelic, et al. 2021). For mouse, these statistics are derived directly from 3D rVANs (Blinder et al. 2013; Gagnon et al. 2015) (**Fig. 1A**). For human, the same statistics are derived from published statistical distributions (**Fig. 1A**).

**Fig. 1:**
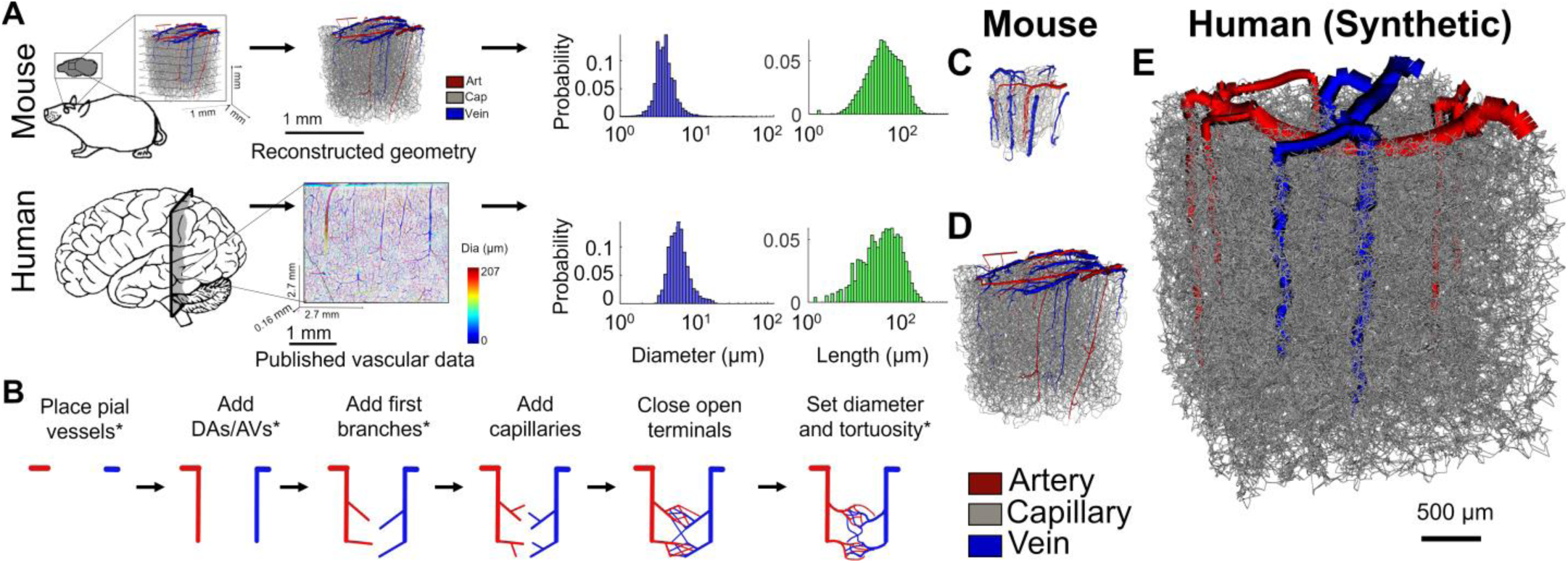
Overview of the iCNS algorithm used for generating synthetic VANs, with geometric and topological features matched to pre-defined statistical distributions, and comparison of mouse reconstructed VANs to a human synthetic VAN. (A) Statistical distributions for mouse vasculature are derived from reconstructed VANs and used to synthesize VAN models specifically for mouse cortex. Statistical distributions for human vasculature are derived from published histology data. The vascular distributions of vessel diameter (blue) and length (green) histograms are used to set the diameter and tortuosity spectra of the synthetic VAN. (B) We synthesize new connections for the VAN using different topological properties for each stage. The final stage adds geometric details to the vessels including tortuosity and diameter. The steps of the iCNS algorithm that were modified in this work are annotated with an asterisk (*). DA = diving arteriole; AV = ascending venule. (C–D) Mouse reconstructed VANs from two laboratories. The 3D renderings of the VANs are colored-coded by anatomical labeling, where red indicates arteries, gray indicates capillaries, and blue indicates veins. (C) Smaller reconstructed VANs derived from *in-vivo* imaging create VANs with 1,300 segments (Gagnon et al. 2015), while (D) larger reconstructed VANs from *ex-vivo* imaging have >14,000 segments (Blinder et al. 2013). There are fewer arteries than veins in these mouse models. (E) In contrast, the synthetic human VAN has 137,500 segments. The human VAN consists of a larger proportion of arteries compared to mouse but a similar proportion of veins. The capillary volume fraction (as a percentage of total vascular volume) is smaller in the human vasculature (44%) than in the mouse (∼60%), however this is not easily seen in the rendering due to the greater overall number of capillaries in the human model.

In order to account for topological differences between humans and mice, we modified four stages of the iCNS algorithm. One key microvascular difference between species is the ratio of diving arterioles to ascending venules (Blinder et al. 2013; Schmid, Barrett, et al. 2017). The targeted ratio was achieved by modifying stages that generate pial and penetrating vessels. We also noted asymmetric branching densities of arteries and veins in the mouse data (see *Results*). We calculate and report the density of these arteriolar and venular branches used in the synthesis in **Table 1**. We note that these branching arterioles and venules are not counted in the commonly reported artery-to-vein ratio (Cassot et al. 2010; Blinder et al. 2013; Duvernoy et al. 1981) and are added symmetrically in prior synthesis algorithms (Linninger et al. 2019; Hartung, Badr, Mihelic, et al. 2021). To incorporate the observed branching densities, we modified the stage that adds the first branches to the diving arterioles and ascending venules.

We also modified the final stage that sets geometric properties of the vessels. The last stage replaces straight segments with tortuous (curved) ones. The previous method employed to achieve this used a Bezier curve approximation (Hartung, Badr, Mihelic, et al. 2021; Linninger et al. 2019) that created unrealistic bifurcations with overlapping vessels and numerical inaccuracies during oxygenation and magnetic field calculations. To increase anatomical realism and reduce vessel overlap, we devised a new approach using a dictionary of reconstructed vessels created from a reconstructed VAN. Each straight vessel segment is then replaced with a real vessel from this dictionary (see *Supplementary Notes* for more details).

#### 3.1.1 Synthesizing and validating VAN models for mouse cortex

We first synthesized five mouse VANs with properties equivalent to those of mouse rVAN1 reconstructed by Kleinfeld and colleagues (Blinder et al. 2013) that was previously used for steady-state hemodynamic and oxygenation simulations (Hartung, Badr, Moeini, et al. 2021; Gould et al. 2017; Linninger et al. 2019). We enforced an arteriole-to-venule ratio of 1:3 for diving and pial vessels for mouse VANs (Schmid, Barrett, et al. 2017; Blinder et al. 2013) as opposed to the previously implemented 1:1 ratio (Hartung, Badr, Mihelic, et al. 2021; Linninger et al. 2019). Moreover, in order to achieve VAN synthesis, vascular structures must not only be imaged and reconstructed in 3D, but a graph representation capturing the connectivity of the network must also be computed. Unlike geometric statistics of vasculature, which are analyzed as frequency of occurrence within a unit volume, vascular synthesis requires statistics of specific properties (e.g., diameter, length, tortuosity) tallied by vessel count (e.g., the number of vessels with a given diameter). Moreover, connectivity patterns between adjacent vessels must be known, which can only be calculated from a network graph or by manually counting the numbers of connections (which is only practically feasible for datasets containing a small number of vessels).

#### 3.1.2 Synthesizing VAN models for human cortex

Our human VAN was synthesized using published geometric and topological statistics (Cassot et al. 2006; 2009; Schmid, Barrett, et al. 2017). **Fig. 1** visualizes the human sVAN compared to mouse rVANs created from *in-vivo* (Gagnon et al. 2015) or *ex-vivo* (Blinder et al. 2013) data. For input parameter values we used published vessel density, diameter, and length distributions (Cassot et al. 2006). For the topology of the VAN, we used an arteriole-to-venule ratio of 2.1:1 (Schmid, Barrett, et al. 2017; Cassot et al. 2009; Lauwers et al. 2008), and enforced the densities of vessel segments branching off of arterioles and venules as measured from histology data (Cassot et al. 2010). Note, all human statistics originated from the same published histology dataset (Duvernoy et al. 1981) as summarized in the *Supplementary Notes*.

### 3.2 Simulating hemodynamics with VAN models

Our dynamic biophysical simulation platform has three distinct steps. The first step calculates the dynamic blood flow responses in the VAN model. The second step simultaneously computes the oxygen distribution via advection through blood, exchange between blood and tissue, and diffusion through and consumption by the tissue. The final step calculates the time-varying T_2_*-weighted signal caused by the variations in deoxyhemoglobin in blood within an applied external magnetic field. For all simulations (humans and mice) we use the same physiological parameter values taken from mouse, for consistency (see **Supplementary Table S4**). This allowed us to investigate the effects of the VAN model (geometry and topology) on the properties of the BOLD response.

#### 3.2.1 Blood flow computations

The active response to neuronal activity originates in the arterioles and propagates upstream through the arterial tree. Our hemodynamic simulations thus use arterial dilations and constrictions recorded in mouse and rat primary somatosensory cortex responding to a 2-s forepaw stimulation (Tian et al. 2010; Uhlirova et al. 2016) as described previously (Gagnon et al. 2015). These recorded diameter changes were assigned as a function of branching level and cortical depth. These arterial diameter changes then induce changes in vascular resistance, pressure distribution, and blood flow as described below.

An initial branching level across vascular hierarchy was assigned in the provided rVAN models, which provided measures of the numbers of first-order branches, second-order branches, and so on. We noted, however, some errors in the branching level assigned to several arteries and precapillary arterioles in the provided rVAN models (which were initially classified using a simple diameter threshold (Blinder et al. 2013)) in which some precapillary arterioles were incorrectly labeled as diving arterioles. This anatomical labeling artifact in which branching level is incorrectly assigned is inconsequential for most VAN applications, however because our inputs are arterial dilations measured in specific branching levels, this must be corrected to properly generate the hemodynamic responses. To ensure consistent branching level assignment across all VAN models and proper assignment of active diameter changes, we algorithmically separated the branching precapillary arterioles from the diving arterioles (see *Supplementary Notes*).

The pressure redistribution after arterial dilation leads to a passive dilation (or “ballooning”) in the downstream capillaries and veins. The relationship between blood pressure and volume is modeled for each blood vessel as previously proposed (Boas et al. 2008) using the relation

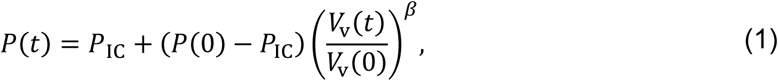

which can be rewritten as

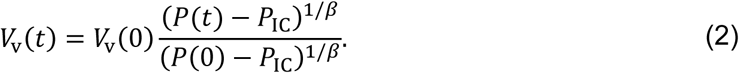

Here, *P*(*t*) is pressure as function of time *t*, *P*_IC_ is the static intracranial pressure, *V*_v_(*t*) is the time-varying vessel volume, and the exponent *β* captures the vessel compliance.

The changes in capillary and venous volume (i.e., their vessel-diameter changes) are calculated by integrating these equations in time, as described previously (Gagnon et al. 2015). This integration is governed by a parameter, *τ*, accounting for the delayed vessel response from wall elasticity and the surrounding tissue viscosity and is often referred to as the “viscoelastic parameter” in the Balloon Model (Buxton et al. 2004; Polimeni and Lewis 2021). The volume changes are then calculated using

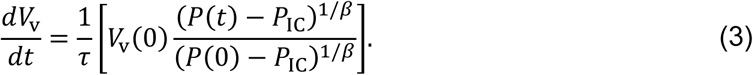

To solve for pressure and flow, we then impose mass conservation and a constitutive Hagen-Poiseuille relationship. We use a timestep of 0.025 s for temporal integration, balancing speed of convergence and stability. The pressure is held constant at the arterial and venous terminals with values listed in **Supplementary Table S4**, although time-varying pressure may be appropriate accounting for whole-brain pressure redistribution in response to localized activation as previously suggested (Pfannmoeller et al. 2021).

#### 3.2.2 Oxygenation calculations

The computational bottleneck in previous VAN-based BOLD simulations was the oxygenation calculation. Previous 3D oxygenation simulation methods use tetrahedral meshes contouring each blood vessel (Fang et al. 2008; Gagnon et al. 2015). These methods, while accurate, suffer from geometrical and topological drawbacks. The numerous tetrahedral elements required to contour small vessels degrade mathematical solvability (a geometrical drawback). Curved blood vessels also require a greater number of small mesh elements to resolve. Furthermore, generic Finite Element Method problems common to non-uniform tetrahedral structures lead to meshes with variable numbers of connections, increasing irregularity in the corresponding system of equations (a topological drawback). These drawbacks increase the size and complexity of the simulation, increasing the runtime and reducing mathematical solvability as explained elsewhere in the context of VAN oxygen simulations (Hartung, Badr, Moeini, et al. 2021). Additionally, previous VAN models use nonlinear neuronal oxygen consumption kinetics (Gagnon et al. 2015). Here, we used a linear kinetic model for efficient computations without sacrificing accuracy, which is described below.

We previously proposed a method that directly couples the 1D graph network representing the VAN structure embedded in a 3D tissue mesh (Hartung, Badr, Moeini, et al. 2021). Unlike more advanced 1D-3D coupling methods (Kuchta et al. 2021), this “1D-3D Finite Volume Method (FVM)” (also known as a “Dual-Mesh” technique (Hartung, Badr, Moeini, et al. 2021)) couples the 1D mesh representing the vascular topology and the 3D mesh representing the surrounding tissue using simple finite differences. We extended this 1D-3D FDM to compute dynamic changes occurring during functional hyperemia. This method couples blood vessels and surrounding tissue by accounting for the proportion of blood and tissue within each regular hexahedral (or “voxel”) mesh element (akin to representing the “partial volume” of each compartment within each mesh element), avoiding the need to contour the mesh to the blood vessels. This offers geometrical and topological advantages. Using hexahedral mesh elements reduces the overall number of elements, and avoiding contouring the blood vessels reduces mesh density (geometric advantages). This also leads to fast convergence and high numerical accuracy even at coarse grid resolutions (Hartung, Badr, Moeini, et al. 2021). The regular grid also has a fixed number of equation variables with flux vectors orthogonal to the cross-section between adjacent mesh elements, providing increased numerical accuracy and stability (topological advantages related to the connectivity of the discrete elements used for computation). These advantages are particularly important because the extravascular mesh accounts for 97% of the oxygenation simulation domain (assuming 3% cerebral blood volume). We chose to impose linearity, further improving solving time and stability. The 1D-3D FDM method is also amenable to coarse-grid initialization strategies that enhance convergence of baseline oxygen profiles used to initialize our hemodynamic simulations.

For dynamics, we added dynamic accumulation (temporal derivative) terms in each mass-balance equation and a numerical integration of temporal derivative terms to the model. We also added an additional hemoglobin-bound oxygen phase as in previous BOLD simulations (Fang et al. 2008; Gagnon et al. 2015) which interacts with blood plasma. Unlike the previous model using the nonlinear steady-state Hill equation, we implemented and validated an equivalently accurate linearized dynamic model (see *Supplementary Notes*).

Our oxygen transport model computes oxygen in three phases: hemoglobin-bound, free plasma, and tissue oxygen. Two phases are carried through the blood vessels: the hemoglobin-bound 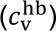 and free plasma 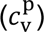 oxygen. The hemoglobin-bound phase is calculated with standard advection-dissociation mechanics given by

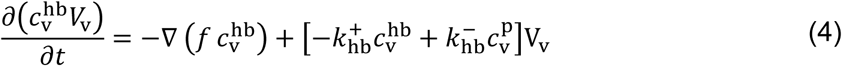

where *f* is volumetric blood flow, 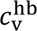 is the concentration of hemoglobin-bound oxygen, 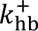 is the oxyhemoglobin dissociation rate, 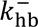 is the hemoglobin-oxygen binding rate, and *V*_v_ is the blood vessel volume. This expression captures how blood flow advects hemoglobin-bound oxygen while dissociating freely into the blood plasma. The plasma-phase is calculated using the relationship

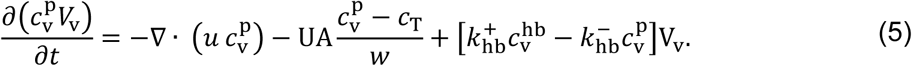

Here, *U* is the endothelial mass transfer coefficient, *A* is the vascular (luminal) cross-sectional area, *c*_T_ is the tissue oxygen concentration, and *w* is the blood vessel wall thickness. This expresses how blood flow advects free oxygen in the plasma while being supplied by oxygen dissociating from the hemoglobin-bound phase. This plasma-phase oxygen then undergoes a diffusive mass transfer across the blood-brain-barrier into the surrounding tissue.

This tissue-oxygen phase diffuses and is metabolized by local neurons and glia. Tissue oxygen is calculated with the expression

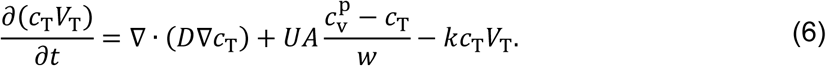

Here, *V*_T_ is the extravascular mesh volume, *D* is oxygen diffusivity, and *k* is the cerebral metabolic rate of oxygen consumption (CMRO_2_). The parameter *k* is estimated from an oxygen extraction fraction (OEF) of 30% and a perfusion rate of 100 ml/100 g/min (Gagnon et al. 2015) with an assumed average pO_2_ of 45 mmHg. We also calculate the equivalent zeroth-order reaction rate constant giving equivalent consumption rate (see *Supplementary Notes*). We held *k* constant in time and space as previously proposed (Gagnon et al. 2015). The tissue observes a no-flux (symmetry) boundary condition on all domain edges. An implicit Euler scheme was used for temporal integration and a finite volume method was used for spatial integration. A fixed boundary condition of 102 mmHg oxygen tension in the plasma and 92% saturation was fixed at arterial inlets. We ensured sufficient mesh resolution at baseline by integrating in time and confirming oxygen changes were <0.00001% in all vessels. We sought the coarsest timestep for computational efficiency but verified the timestep was sufficiently small by simulating with a 10× finer timestep and then ensuring oxygen changed by <0.1% in all vessels.

#### 3.2.3 MRI signal calculations

To simulate hemodynamic changes impacting the BOLD fMRI signal, we calculate water protons in the extravascular domain dephasing from local magnetic field inhomogeneities. These inhomogeneities caused by magnetic susceptibility differences (when placed in an external magnetic field, *B*_0_) between cortical gray matter and paramagnetic deoxyhemoglobin as computed using a finite perturber method (Pathak et al. 2008; Koch et al. 2006) as previously described (Gagnon et al. 2015; Pfannmoeller et al. 2020). For completeness, we summarize these computations here.

First, the VAN anatomy is mapped onto a regular grid (a new grid is calculated for every time step, accounting for CBV changes) and labeled with corresponding oxygen saturation (*S*). The magnetic field inhomogeneities (Δ*B*(*x*, *y*, *z*)) are then computed by convolving the magnetic susceptibility map (Δ*χ*)

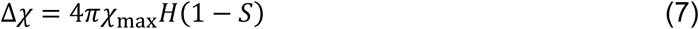

with a convolution kernel (*K*)

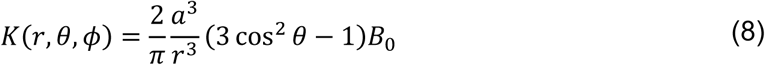

representing field offsets generated by a susceptibility point source. This convolution in the spatial domain can be computed as a multiplication in the Fourier domain as

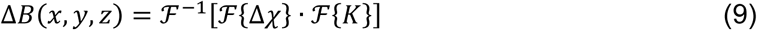

as described previously (Pathak et al. 2008). Here, *a* is the grid spacing, *r* is the distance between the kernel center and each grid point, ℱ is the Fourier operator, *H* is the local hematocrit value, and *χ*_max_ is the susceptibility difference between fully oxygenated and fully deoxygenated blood. Variable *θ* is the alignment angle between the main MRI magnetic field and the vector formed between the kernel center and each grid element. The magnetic moment is assumed uniform across all azimuthal angles *φ*.

Our simulated MRI protocol used a standard gradient-echo (GE) pulse sequence to measure the T_2_*-weighted BOLD signal. We use a magnetic field strength of 7T, a 90° excitation flip angle, and a frequency-encoding gradient in the *x*-direction as listed in **Supplementary Table S4**. Our frequency-encoding gradient pulse sequence uses a pre-phaser of 10 ms applied 10 ms after the time of excitation (*t*=0 s). We then immediately begin a 20-ms readout gradient to generate a gradient-echo at 30 ms.

Randomly placed protons then diffuse by random walk through the extravascular domain. The radiofrequency (RF) excitation pulse (at *t*=0 s) phase-aligns all protons. Between *t*=0 s and *t*=TE, the proton phase (*φ*_ev_) is updated according to the local magnetic field by

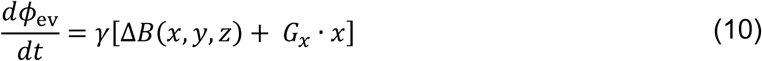

accounting for the field inhomogeneities (Δ*B* and *G_x_*). Here, *G_x_* is the frequency-encoding magnetic field gradient used for image encoding, x is the x-coordinate of the proton, and *γ* is the gyromagnetic ratio of water protons. We include readout image-encoding gradients to ensure our results are comparable to previous simulations (Gagnon et al. 2015); although this gradient introduces a small amount of inadvertent diffusion weighting into the MR signal and subtly impacts the BOLD effect by partially adding to or counteracting extravascular gradients around large veins (Berman et al. 2021). The resulting voxel signal (*S*_GE_) is the mean of all complex-valued signals from individual protons where the signal magnitude reflects phase cancellation, given by

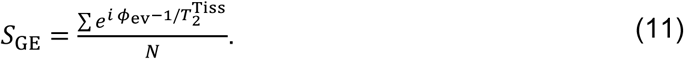

This expression also accounts for the real-valued signal decay associated with irreversible transverse relaxation. Here, *i* is the imaginary unit, 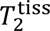 is the transverse relaxation rate for gray matter, and *N* is the total number of protons. We use a grid spacing of 1 μm and a 200-µs timestep as validated in previous work (Gagnon et al. 2015; Pfannmoeller et al. 2020) and compute the complex-valued BOLD response at 0.5-s intervals. We then compute the magnitude of the BOLD signal and report the BOLD response amplitude as the percent signal change relative to baseline using the conventional definition (e.g., *S* = Δ*S*/*S*_0_).For computational efficiency and to reduce memory load for the larger human VAN, all magnetic field calculations were performed with single-precision.

### 3.3 Model parameter value calibration

Our focus in this study was to emphasize the differences in vascular structure on the hemodynamic response. To minimize bias (and the potential for overfitting) due to an arbitrary selection of input parameter values, our simulation inputs were derived from first-principles with only two simple calibrations: (i) the targeted baseline cerebral blood flow (CBF, 100 mL/100 g/min) was achieved by adjusting blood viscosity; and (ii) the targeted baseline oxygen extraction fraction (OEF, 30%) was achieved by adjusting the oxygen metabolism rate coefficient. In both cases, the targeted values for CBF and OEF are well established and taken from the literature.

### 3.4 Evaluating sVAN validity with dynamic BOLD simulations

The geometry as well as baseline simulations of blood flow and oxygenation in our synthetic VANs were previously validated (Hartung, Badr, Moeini, et al. 2021; Hartung, Badr, Mihelic, et al. 2021). To further validate sVANs for dynamic BOLD simulations, we compared the amplitudes and temporal features of the BOLD fMRI responses from five mouse sVANs to those of two rVANs (rVAN1 and rVAN2, providing two independent reconstructions from the same tissue preparation and methods (Blinder et al. 2013)). Because a human 3D rVAN does not exist, we qualitatively compared human sVAN BOLD simulation results to published fMRI responses. We compare peak amplitude (in units of percent signal change), amplitude of the post-stimulus undershoot, rise-time, and fall-time of the main BOLD peak.

### 3.5 Extrapolating arterial responses to the human VAN

Equivalent *in-vivo* single-vessel data that represent the vascular response to neuronal activation in mouse VANs (Uhlirova et al. 2016; Tian et al. 2010) cannot be imaged in humans noninvasively. Therefore, we extrapolated (or scaled) the measurements from the ∼1-mm-thick mouse cortex to our 2.5-mm-thick human cortex model. Our predicted dilation schemes incorporate the well-known upstream dilation propagation from the smallest arterioles up the vascular hierarchy to the arteries on the pial surface. We model the coordinated vascular response to neuronal activity using progressively complex extrapolation schemes.

Scheme 1 (**Fig. 2A**) directly assigns the recorded arterial dilations as a function of relative cortical depth, i.e., each time-course is assigned to the same relative depth in all VANs as a percentage of cortical thickness. This Scheme theoretically assigns dilations to equivalent vascular hierarchical levels between species. This simplest model, Scheme 1 is somewhat unrealistic yet provides a reference “null” model.

**Fig. 2:**
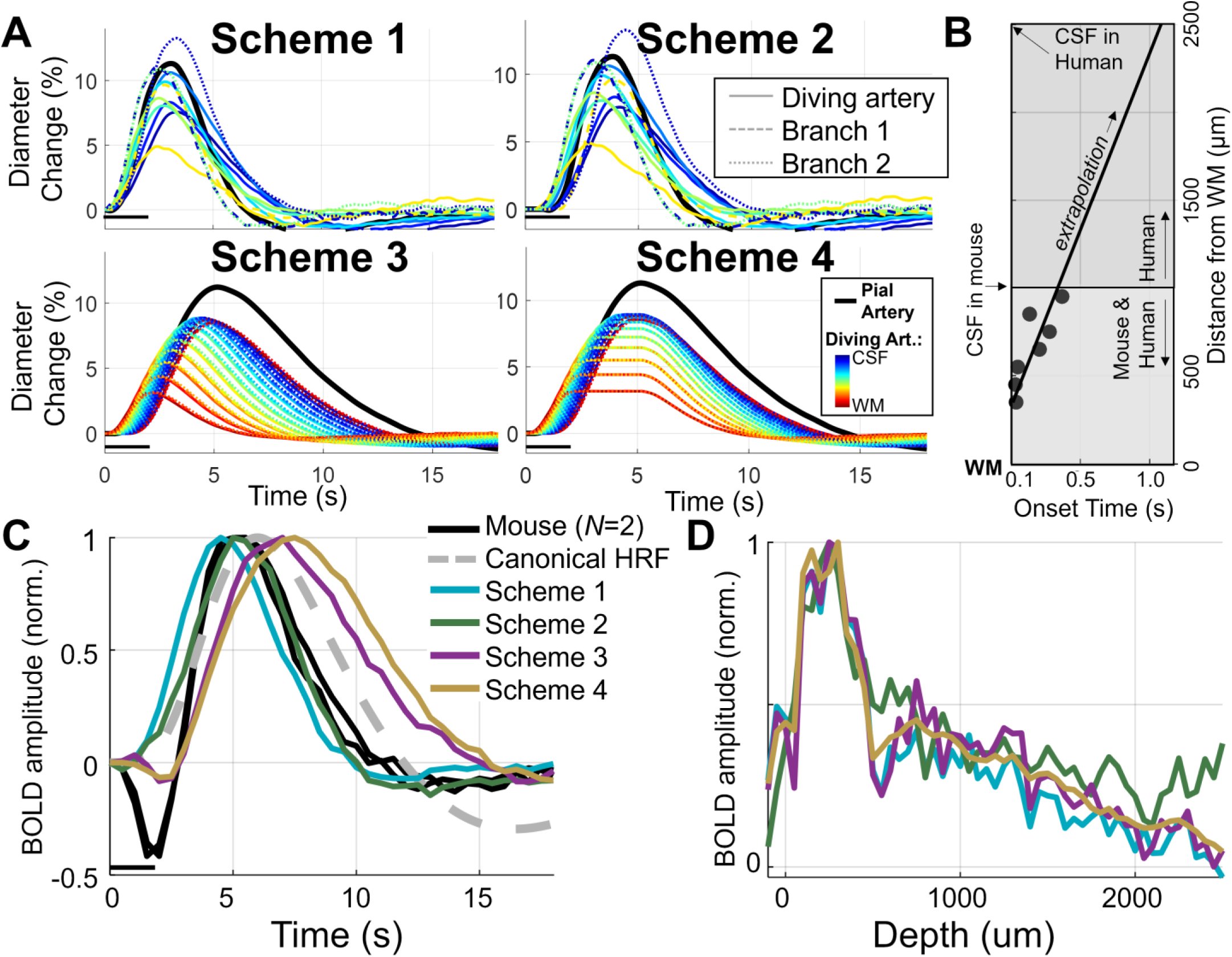
Comparison of resulting arterial dilation inputs using different Schemes for scaling arteriolar dilations from rodent to human and resulting BOLD responses from mouse and human VANs. (A) Time courses of our four dilations Schemes, with the sub-plots each corresponding to a different dilation Scheme. We report the pial arteries (black line), the diving arteries (DA, colored solid lines), branch-1 arterioles (dashed lines), and branch-2 arterioles (dotted lines). The diving arteries and branching arterioles are assigned different dilation time courses according to their cortical depth. The color coding for the diving arteries and branching arterioles corresponds to their respective cortical depths as indicated in the legend, with dark blue corresponding to the cerebrospinal fluid (CSF) interface at the pial surface and dark red corresponding to the white matter (WM) interface. (B) An example of a linear fit to one example timing parameter (onset time) used to extrapolate the mouse vessel diameter recordings for the human VAN simulations. The data between the WM and CSF interface in mouse (circles) were fit with linear regression (solid line) that was then extrapolated to the thicker human cortex. Note vessel segments in the lowest depths are assumed to have a synchronous onset at time *t*=0 s). (C) The BOLD responses from the different Schemes are plotted. The black traces represent two BOLD responses simulated from mouse rVAN1 and rVAN2. The BOLD response from the human sVAN using Scheme 1, 2, 3, and 4 are shown in blue, green, purple, and gold, respectively. Here we compare only the timing between mouse and human simulations therefore the amplitudes of the BOLD responses have been normalized to their respective peaks. We also show the stimulus duration (black bar). For reference we added the default canonical hemodynamic response function (HRF) for humans provided by the SPM analysis package (Ashburner 2012). (D) Cortical depth-profiles of BOLD responses under Schemes 1, 2, 3, and 4 (blue, green, purple, and gold lines respectively). All Schemes exhibit the expected peak BOLD response at the pial surface due to the large pial vein in the human VAN.

Scheme 2 assumes that the dilation signal propagates upstream at the same velocity in both species. We calculated the speed of the upstream propagation of dilation by calculating the onset timing of each measured dilation trace at each cortical depth using a threshold of 0.2% relative change. We then performed linear regression on these onset times as a function of cortical depth. In these data, the deepest depth (near to the white matter) corresponds to the shortest onset time. The slope of this regression yielded the propagation speed (1.96 mm/s). The relative depth assignment matches that of Scheme 1 however the onset time of each time-course is adjusted according to absolute cortical depth (see *Supplementary Notes*).

Scheme 3 varies the four parameters that most clearly vary with depth in the rodent data: (i) onset time, (ii) rise-time, (iii) fall-time, and (iv) amplitude. Parameters (i)–(iii) seemingly vary linearly with depth (see *Supplementary Notes*). Parameter (iv) is estimated using a quadratic polynomial chosen as the simplest model once a linear relationship was ruled out. This second-order polynomial trend in amplitude is physiologically plausible as vascular responses may peak in central cortical depths (e.g., Layer IV) where neuronal activity is often strongest. The linear fit for the remaining parameters was chosen to avoid overfitting. We discuss the impact of this assumption in the *Discussion*.

In Scheme 3, the deepest vessels return to their baseline diameter before the pial vessels reach their peak dilation. It is unlikely that the large upstream feeding arteries would be dilated when the small downstream arterioles are not (Uhlirova et al. 2016). To rectify this, we adapted Scheme 3 slightly to create Scheme 4, where we sustained peak dilation in these deepest arterioles until all active vessels are fully dilated. Once all active segments have reached their maximum dilation, the return to baseline begins in all active segments simultaneously.

### 3.6 Evaluating hemodynamic differences between mouse and human

To evaluate potential impacts of vascular differences between mice and humans, we compared the hemodynamic responses to long-duration stimuli. Passive venous dilation was predicted to be more pronounced during long-duration stimulation compared to short-duration (Drew et al. 2011; Polimeni and Lewis 2021; Havlicek and Uludağ 2020), as the longer stimulus time presumably should enable full venous dilation. Due to the longer paths and thus greater absolute distances (in units of mm) between the sparser arteries and veins in humans, we hypothesized that this effect would be more pronounced in the human VAN than the mouse VAN. To test this, we began with a short-duration stimulus using the measured arterial dilation recordings for the mouse rVAN (Uhlirova et al. 2016; Tian et al. 2010) and the Scheme 3 estimation for the human sVAN. We chose Scheme 3 because it is the simplest extrapolation that resulted in realistic BOLD timing for the human sVAN (see *Results*). To create the active dilations for the long-duration stimulus, we sustained the peak dilations of the short-duration stimulus for an additional 18 s to create a 20-s total stimulation duration. We also adapted the viscoelastic parameter (*τ*) as previously suggested (Buxton et al. 2004; Polimeni and Lewis 2021; Havlicek et al. 2017) to account for the different levels of venous ballooning for short- and long-duration stimuli (Hillman et al. 2007).

### 3.7 Investigating contributions to BOLD from individual compartments

As our realistic VAN modeling approach enables simulating BOLD responses within arbitrarily small MRI voxels across the microvascular hierarchy, we simulated cortical-depth or “laminar” BOLD response profiles corresponding to several microanatomical variations. To generate these cortical-depth profiles, we simply binned the BOLD response amplitude into equally-spaced depths (50 μm thickness) for each VAN model. We compared the laminar BOLD response profile for rVAN1 and rVAN2, by simulating BOLD responses from all vessels and also for each compartment. We achieved this compartment-specific simulation and analysis by first simulating the hemodynamic response for the entire vasculature (which we term the “full response”). Then, we performed a second simulation with a constraint placed on the hemodynamics: we only allow a single vascular compartment to vary in diameter and oxygen content (extracted from the “full response”, to ensure realism), while the other vessels remain at baseline throughout the simulation. We term this the “constrained response”.

Our reconstructed VAN models are derived from cortical locations away from large pial veins, thus they do not reflect the typical peak BOLD amplitude at the pial surface. To investigate the impact of large pial veins on the laminar BOLD profile, we conducted additional simulations with a large vein with radius 75 µm grafted to the model (see **Fig. 6B**) at the pial surface. We assigned oxygenation and dilation time-courses to this grafted vein using the average of the oxygenation and dilation dynamics from all pial veins in the original, unaltered VAN simulation.

## 4 Results

### 4.1 VAN model synthesis

Our first step towards human BOLD simulations was VAN synthesis (see *Methods*). The synthetic human VAN shown in **Fig. 1** is substantially larger than mouse VANs in dimensions and overall numbers of vessels.

### 4.2 Validation blood and tissue oxygen computations

To confirm the validity of our new dynamic oxygenation model (see *Methods*), we compared our predicted intravascular oxygen tension to both that of an existing model (Gagnon et al. 2015; Fang et al. 2008) and *in-vivo* measurements (Gagnon et al. 2015). The baseline oxygenation, sorted by vessel diameter, exhibited close agreement between all three, as shown in **Fig. 3A**.

**Fig. 3:**
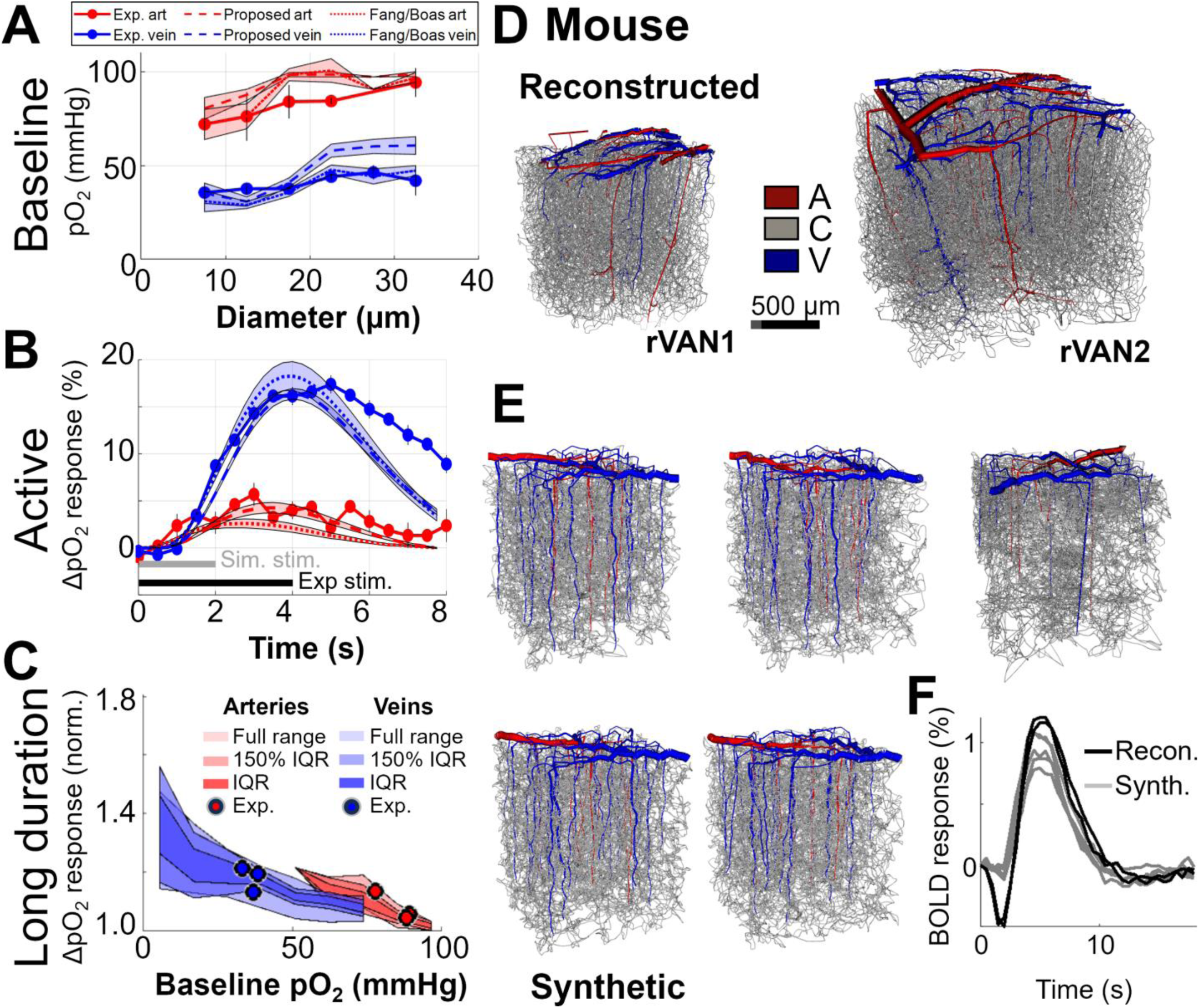
Validation of oxygenation at baseline and activation and BOLD responses in synthetic VAN structures. (A) We overlaid the baseline oxygen tension in arteries and veins for three datasets binned by diameter. We show experimentally observed tension (solid lines), simulated tension using the previous gold-standard method for similar simulations, termed the “Fang/Boas” method (dotted lines) and using our proposed method (dashed lines). The results are plotted separately for arteries (red) and veins (blue). (B) We also compared the active responses with experimental data (Yaseen et al. 2011). The annotation bars indicate the stimulus duration for the experiment (black, 4-s-long stimulation) and simulations (gray, 2-s-long stimulation). (C) A second such comparison to data from another study (Vazquez et al. 2010), in this case with a longer-duration stimulation. Here, arteries and veins are indicated by red and blue, respectively. The light shading indicates the full range of simulated values, the medium shading indicates 150% of the inter-quartile range (IQR), and the dark shading indicates the IQR. The circles indicate the measured data. (D) Two example reconstructed VANs (rVANs) provide reference simulated BOLD responses. (E) Five examples of synthetic VANs (sVANs) were generated from the statistics of rVAN1. The sVANs exhibit qualitative similarity (similar amplitude, rise-time, fall-time, and post-stimulus undershoot magnitude) to the reconstructed VANs from which they were derived. (F) BOLD responses for the two reconstructed and five sVANs show comparable amplitudes and time-courses. The initial dip of the BOLD response is more prominent in simulations based on these rVANs.

We also compared simulations and measurements of dynamic oxygenation responses in mouse somatosensory cortex to forepaw stimulation. We compared simulated responses to 2-s-long stimulation (Tian et al. 2010; Uhlirova et al. 2016) to measurements using a 4-s-long stimulation (Yaseen et al. 2011), shown in **Fig. 3B**. While comparing identical stimuli would be ideal, our simulation requires dilation measurements that are only available for a 2-s-long stimulation, whereas measurements of dynamic intravascular oxygenation responses are only available for a 4-s-long stimulation. Despite subtle differences in stimulus duration and response timing, we observed comparable response amplitudes between the measured responses and the simulated responses. While the simulated responses were smaller and returned to baseline earlier, simulations using both methods are in good agreement with the measurements. The measured responses plateau at roughly the same time as the simulated response peak with both methods, possibly contributing to the agreement despite the different stimulus timing. The proposed method yielded an oxygenation response peak in arteries at ∼5 s (after the stimulus offset), only ∼2 s after the arterial dilation measurements peak. In both experimental and simulated data, the venous oxygenation response is ∼1 s delayed relative to the arterial response. The arterial response, driven by arterial dilations and transit time through the arterioles, is subtly slower when computed with the proposed method than with the Fang/Boas model. Extended comparisons are provided in the *Supplementary Notes*. Our first-order reaction rate model of oxygen consumption resulted in an increased metabolic rate of ∼12% during activation (see *Supplementary Notes*).

In order to validate the oxygenation changes during activation, we compared the simulated oxygenation responses to long-duration stimuli in arteries and veins to measured data (Vazquez et al. 2010) (see **Fig. 3C**). Because the arteries and veins in the measured data are larger than those within our VAN models, a comparison of measured oxygen tension across vessels with a wide range of diameters was not possible. (If larger VAN models were available that encompassed a larger volume of cerebral cortex and contained a broader range of vessel diameters including large-diameter vessels, a more direct comparison of simulation and measurement would be possible.) We therefore validated our simulated changes in oxygen tension during activation by comparing values in large arterioles and venules against corresponding values from the empirical data. We find that within the arteries and veins (in this case from rVAN1) the empirical data matches well with our simulations. We note that many factors influence the BOLD response amplitude (e.g., stimulus duration), however these BOLD response amplitudes are within ranges of published literature for similar 2-s stimulus duration at 7T; see **Supplementary Table S2** for a summary of expected values of this and other physiological parameters.

### 4.3 BOLD simulations of synthetic VANs

Because this is the initial application of synthetic VANs (sVANs) produced by the iCNS algorithm to dynamic BOLD response simulations, we first evaluated our sVANs as a valid substitute for reconstructed VANs. Two rVANs (**Fig. 3D**) generated from mouse somatosensory cortex (Blinder et al. 2013) were simulated (see *Methods*) to determine the amplitude and timing of the expected BOLD response. We compared these responses to those from five sVANs (**Fig. 3E**) statistically matching rVAN1. Note, these VANs span the entire cortical thickness in mouse (1–1.4 mm) (Blinder et al. 2013), substantially larger than previous dynamic BOLD simulations (Gagnon et al. 2015; Genois et al. 2021; Pfannmoeller et al. 2020; 2021). The sVANs consistently elicited BOLD amplitudes and temporal features comparable to rVANs (**Fig. 3F**). This consistency confirms the similarity between BOLD responses of sVANs and rVANs, validating our sVANs for such simulations. We note that the BOLD “initial dip” is more pronounced in rVANs (see *Discussion* and *Supplementary Notes*).

Our blood flow and oxygenation simulations required <1.5 CPU hours, and BOLD simulations added <1.5 CPU hours per VAN. This represents a ∼570× speed-up (primarily due to the improved oxygenation model) compared to the previous simulation framework without sacrificing accuracy.

### 4.4 Influence of vascular branching patterns on the BOLD response amplitude

We observed the strongest BOLD responses in sVANs with relatively few branches off diving arterioles and many branches off ascending venules. Upon further investigation, this asymmetric branching pattern was also detected in the rVANs (see **Fig. 4** and *Supplementary Notes*). We term this feature the minimal arterial branching (or “MAB”) property. We confirmed this MAB property first in mouse rVANs, with 5.0 and 12.5 branches/mm from diving arterioles and ascending venules, respectively, a 1:2.5 ratio. We then calculated a similar branching asymmetry by analyzing published histology data (Cassot et al. 2010) of human cortical microvasculature (Duvernoy et al. 1981) with densities of 4.3 and 13.1 branches/mm from the diving arterioles and ascending venules, respectively (a *branching asymmetry ratio* of 1:3). Investigating any related impact on capillary asymmetry, however, is outside the scope of this work.

**Fig. 4:**
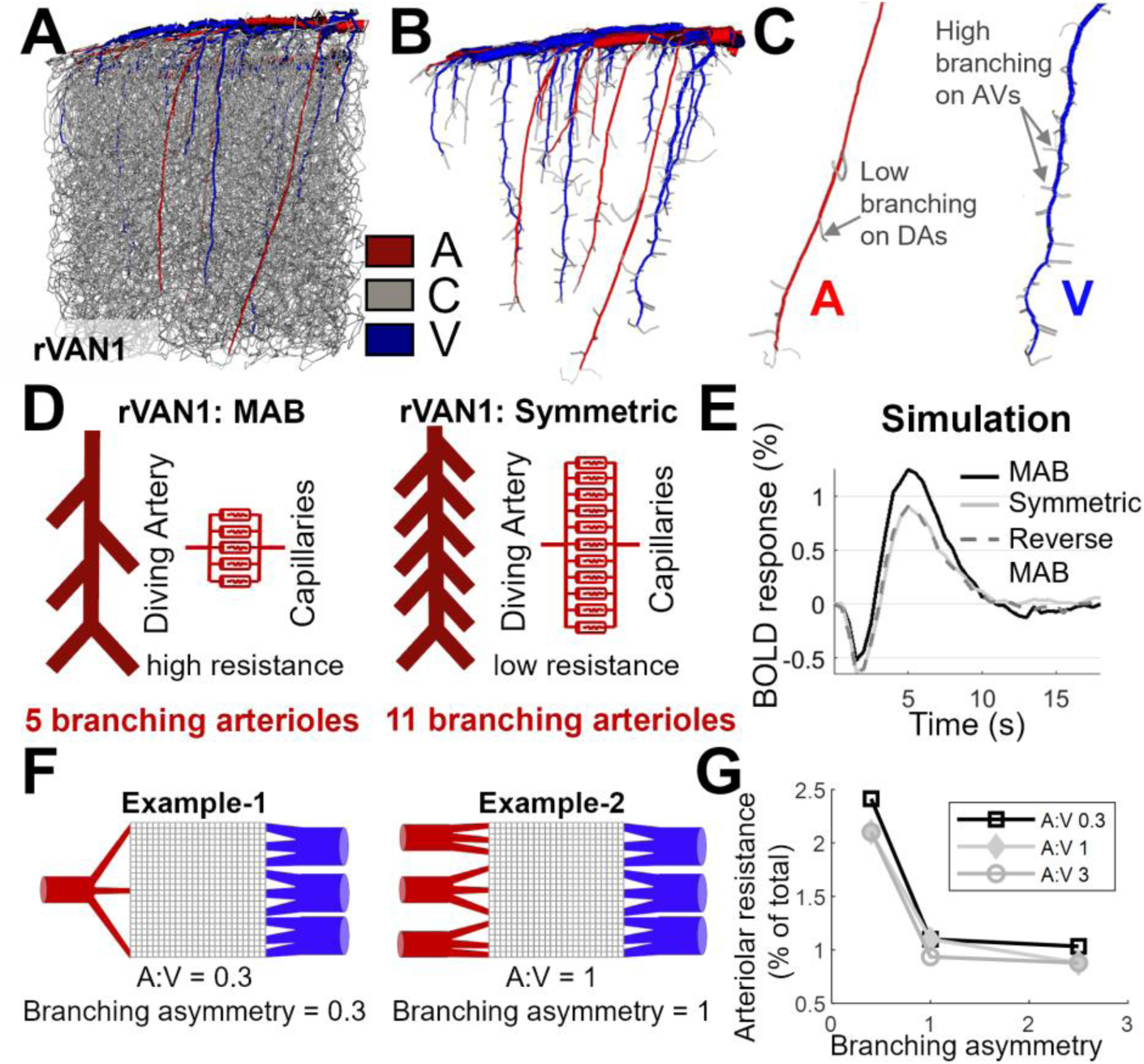
Mouse cortical microvasculature exhibits asymmetric branching between diving arteries and ascending venules. (A) A rendering of rVAN1 where arteries are labeled in red, veins in blue, and all other segments (arterioles, capillaries, and venules) are labeled gray. (B) rVAN1 after removing the capillaries better visualizes the branching patterns of the diving arteries and ascending venules. (C) A representative diving artery and ascending vein further highlights the asymmetry in the branching density between arteries and veins. (D) Schematic representation of diving arterial branching patterns from two hypothetical rVANs with differing numbers of branching arterioles and their corresponding resistors based on a circuity-theory analogy. Here, rVAN1 MAB has fewer branching arterioles per diving artery (5 pictured here) and rVAN1 Symmetric has more branching arterioles (11 pictured here). (E) The BOLD response simulations from rVAN1 a small number of branching arterioles (i.e., with MAB, solid black line), rVAN1 with symmetric branching (solid gray line) and reverse MAB (dashed gray line), both having a large number of branching arterioles (i.e., without MAB) result in differing BOLD response amplitudes. The simulations based on rVAN1 with symmetric branching and reverse MAB both exhibited a stronger initial dip and a reduction amplitude of the peak BOLD response by greater than 50%. (F) Two examples of a simplified resistor circuit used to investigate the effects of the MAB principle—Example-1 with an A:V ratio of 0.3 (1:3) and branching asymmetry of 0.4, similar to a mouse VAN with the MAB principle, and Example-2 with A:V and branching asymmetry of 1 and 1, respectively. (G) The fractional arteriolar resistance of these branching arteries as a function of branching asymmetry (horizontal axis) and A:V ratio (different traces).

An example rVAN1 is shown in **Fig. 4A**, with its diving arterioles and ascending venules highlighted in **Fig. 4B** and the branches off one representative diving arteriole and ascending venule highlighted in **Fig. 4C**. This example demonstrates the smaller numbers of branches off diving arterioles compared to ascending venules.

The influence of the MAB property on hemodynamics can be understood with a circuit-theory analogy (**Fig. 4D**): dilations during activation lower arterial resistance, and a substantial resistance change is needed to elicit a strong flow response. Therefore, a small number of branches off diving arterioles is favorable because it generates high baseline resistance that can be substantially lowered with activation. In contrast, having more branches is detrimental because it generates low baseline resistance.

Furthermore, to verify the effect of MAB on the hemodynamic response, we performed additional tests using edited rVAN models in which we manipulated the relative numbers of arteriolar and venular branches. We generated two edited rVAN models: (i) an edited version of rVAN1 with symmetric branching, i.e., equal numbers of arteriolar branches and venular branches by adding arterioles connecting the diving arteries to the nearby capillaries; and (ii) an edited version of rVAN1 with asymmetric branching that is the reverse of the observed MAB principle, i.e., more arterial branches than venular branches, by removing venules connecting the ascending veins to the capillary bed while ensuring no capillaries were orphaned or terminated. These steps are detailed further in the *Supplementary Notes* and visualized in **Supplementary Fig. S13**. The resulting BOLD responses generated using these two edited versions of rVAN1 are shown in **Fig. 4E**. As predicted, these results demonstrate a weaker hemodynamic response in rVANs that do not exhibit the MAB principle.

To quantitatively investigate the impact of these branching ratios to the number of feeding arteries and veins, we developed a simple resistor model that allowed us vary the connectivity (i.e., the amount of branching) and total number of arteries, arterioles, capillaries, venules and veins to demonstrate how differences in the artery-to-vein ratio and arteriolar branching affects the hemodynamics in the vascular network. Here we represent the capillary mesh as a square grid for simplicity. **Fig. 4F** includes two example models: Example-1 with 1 diving artery and 3 branching arterioles, and Example-2 with 3 diving arteries and 9 branching arterioles. (Both resistor models shown here have 3 ascending veins and 9 branching venules.)

As a measure of the control the branching arterioles have on blood flow regulation, we quantified the resistance across these branching arteriolar segments as a percentage of the total resistance across the model. Using this measure, more arteriolar resistance translates to greater control over the hemodynamic response by these vessels and, likewise, a lower resistance translates to less control. We evaluated this fractional arteriolar resistance as a function of the intracortical artery-to-vein (A:V) ratio and the branching asymmetry (arterioles:venules). We observed a drop in the arteriolar resistance as the A:V ratio increases—i.e., resistance decreases as more arteries are added, as seen in **Fig. 4G**, as expected. However, we observed a stronger effect of the number of arteriolar branches on the fractional arteriolar resistance—less arteriolar resistance as more arterioles are added, because each new arteriolar branch adds a resistor in parallel to the other branches. This indicates a greater influence of the number of arterioles (which impacts branching asymmetry) on blood flow regulation than the number of intracortical arteries (which impacts A:V ratio).

Because we find this MAB property in both mouse (Gagnon et al. 2015; Blinder et al. 2013) and human data (Cassot et al. 2010) (an exceptional and notable similarity between the species; see *Discussion*), and because VANs with the MAB property generate more realistic BOLD peak amplitudes, we enforced the MAB property in all other sVANs reported in this work.

### 4.5 Simulations of BOLD response dynamics using the human VAN

The human VAN blood flow and oxygenation simulations required 21 CPU hours (an estimated ∼4,600× efficiency increase relative to previous frameworks). We simulated all candidate human dilation Schemes (see *Methods*), each resulting in distinct BOLD dynamics (**Fig. 2**), to determine which response best matched the human fMRI data. All Schemes were extrapolated from existing rodent single-vessel microscopy data. The simulated response using Scheme 1 (the rodent measurements assigned by relative cortical depth) had a faster rise-time to that of the mouse, peaking at 4.5 s but with a smaller initial dip, post-stimulus undershoot, and faster fall-time. Scheme 2, with extrapolated dilation onset timing, was slower, peaking at the same time as the mouse responses, at 5 s, and exhibiting a larger initial dip and post-stimulus undershoot than produced by Scheme 1. Scheme 3, using a four-parameter extrapolation function, generated slower responses than those of Schemes 1 and 2, exhibiting a peak around 7 s, a larger initial dip and considerably delayed PSU. Scheme 4, a copy of Scheme 3 with prolonged dilation in deeper cortical depths, yielded the slowest response of all Schemes yielding a similar response timing as Scheme 3 but with a response peak at 7.5 s. Overall, only Schemes 3 and 4 yielded BOLD response timings similar to those of human fMRI data (Buxton et al. 2004; Lambers et al. 2020). This suggests the larger scale and longer distances traveled by upstream dilation propagation (a geometric feature) may influence the hemodynamic response timing (see *Discussion*). We also note that the cortical-depth profile of the BOLD response matches best with prior observations (based on mouse and human fMRI data) when using Scheme 1, 3, and 4, which exhibit the expected decrease in amplitude from the CSF to the WM interface. However, all Schemes show a peak at the pial surface, ostensibly from the large pial veins.

### 4.6 Predicted differences in hemodynamics between humans and mice

Next, we tested whether differences in vasculature, such as different artery-to-vein ratios, between humans and mice impact the hemodynamic response. Because this ratio is approximately flipped between mice and humans (1:3 in mice and 2.1:1 in humans), we reasoned that the larger arterial blood pool flowing into a smaller number of veins in human compared to mice may cause greater passive compliance, or “ballooning”, in capillaries and veins. Moreover, we anticipated that the longer path length between arteries and veins in humans (caused by the lower pial vessel density) would require a longer activation duration to reach the full dilation in distal capillaries and veins. Thus, we anticipated this passive dilation would be maximal during long-duration stimuli in agreement with predictions that venous ballooning effects are more pronounced for longer-duration stimuli (Hillman et al. 2007; Polimeni and Lewis 2021; Chen and Pike 2009; Mandeville et al. 1999).

We tested this hypothesis with simulations of long-duration (20-s) stimuli. The simulated blood volume responses in each vascular compartment in mouse (rVAN1) and the human sVAN are shown in **Fig. 5A**. Arterial volume differences were expected to reflect different input dilations for each species. Contrary to our expectation, the passive downstream responses were smaller in human than in mouse, with negligible capillary and venous blood volume changes in the human VAN. Note, the post-stimulus undershoot observed in our BOLD simulations ostensibly reflects the post-stimulus constriction in arteries.

**Fig. 5:**
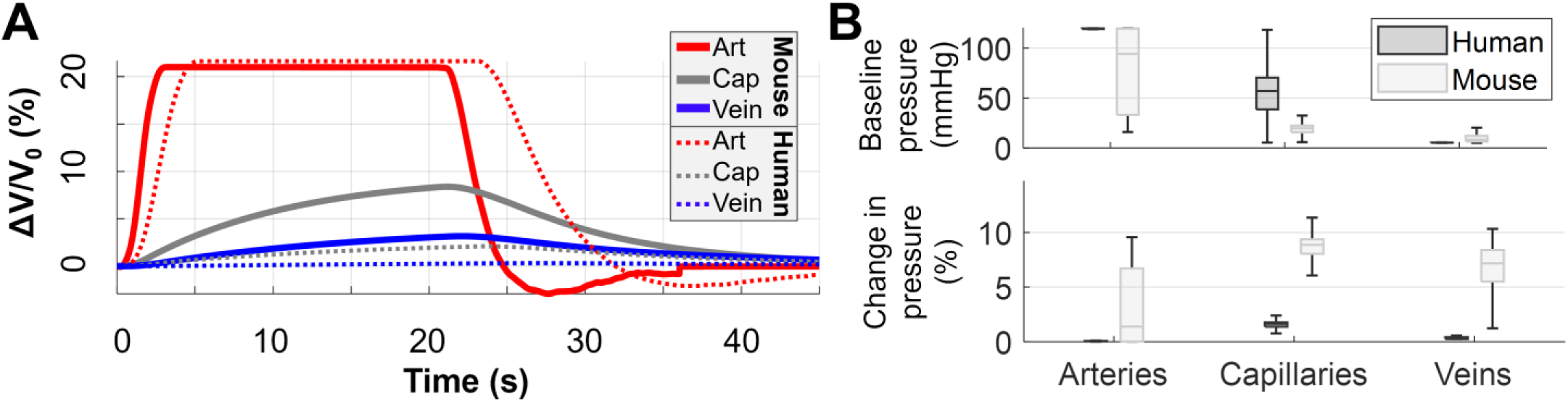
Hemodynamic differences between mouse and human VAN simulations. (A) Blood volume responses (ΔV/V_0_) to long-duration stimulus in a reconstructed mouse VAN and human sVAN shown for each vascular compartment (arteries in red, capillaries in gray, and veins in blue). The black bar indicates stimulus duration. The mouse simulations (thick lines) reflect larger passive ballooning in capillary and venous compartments compared to the human response (thin lines). The human simulations also resulted in slower ballooning compared to the mouse. (B) The baseline pressure distribution in human (dark gray) is plotted against that of mouse (light gray). The changes are broken down by compartment (arteries, capillaries and veins) for comparison. The boxplot indicates the median pressure (horizontal line), the box spans the lower- to the upper-quartile edges and the whiskers extend to 150% of the interquartile range. Outliers are omitted for clarity. The arterial pressure range in the mouse is large, indicating high resistance, while in the human most of the pressure drop is across the capillary bed (arteries and veins are mostly at terminal pressure). The change in pressure due to active and passive dilations in the VAN during activation is also shown for human and mouse. In all compartments, the pressure change in the human VAN is smaller than that in the mouse VAN.

**Fig. 6:**
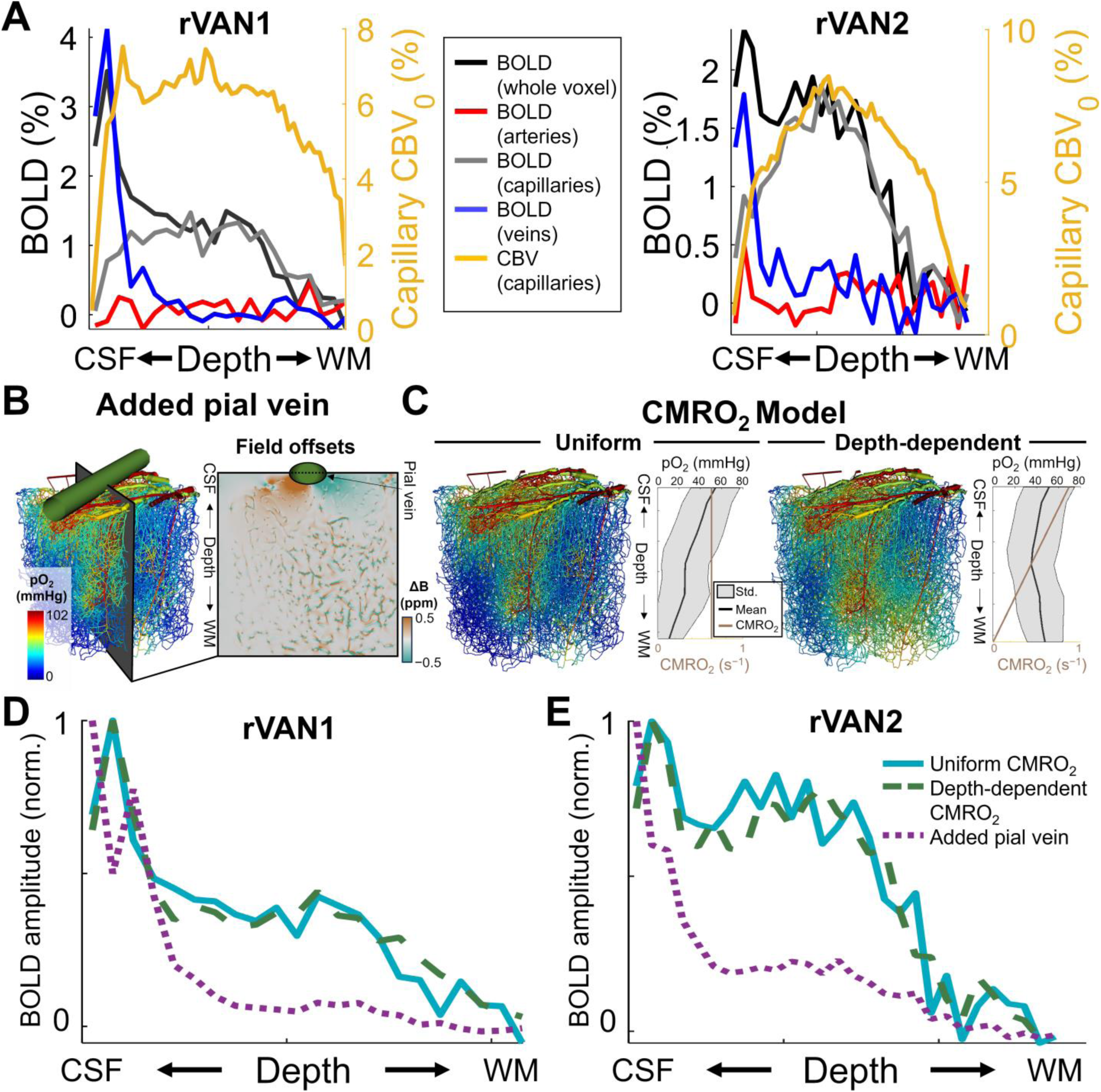
BOLD fMRI responses from individual vascular compartments and with varying geometry. (A) The baseline blood volume of the capillaries (cCBV_0_, gold line) is plotted for rVAN1 and rVAN2 across cortical depth alongside the corresponding depth-dependent BOLD profiles for all vessels, and for the arteries, capillaries, and veins (black, red, gray and blue lines, respectively). The BOLD response profiles from all vessels and from capillaries resemble that of cCBV_0_, with a clear peak in the middle cortical depths. (B) To test for the effects of large-sized veins seen in some locations in mouse cortex and in human cortex, we added a large pial vein to the VAN model. The effect of this large vein is seen in resulting magnetic field offsets, with a large magnetic field at the CSF interface falling off rapidly with distance. (C) Uniform CMRO_2_ across depths leads to low oxygenation (pO_2_) in the lower layers, while a linearly-increasing CMRO_2_ towards the white matter results in a more flat pO_2_ profile across cortical depth. The BOLD responses generated from both rVAN1 (D) and rVAN2 (E) reflect a more gradual falloff between the CSF and WM interface when using linearly-increasing CMRO_2_ towards the white matter compared to uniform CMRO_2_. The large pial vein increases the BOLD response amplitude at the CSF interface, however it has little effect on the BOLD response amplitude for the majority of cortical depths.

To better understand these passive ballooning differences between species, we also compared the baseline hemodynamic states. Using pressure as an indicator of baseline resistance, we compared the histograms of baseline pressure in each vascular compartment. The pressure changes in the arterial, capillary, and venous compartments (**Fig. 5B**) are indicative of the smaller dilations in capillaries and veins in the human VAN. These smaller changes are associated with longer path-lengths and higher effective resistance of the capillary bed compared to mouse VANs (see *Discussion*). This is another way in which distances or geometric features of the human VAN may lead to differences in the hemodynamic response (see *Discussion*). We note that the magnitude of venous dilation is affected by the parameter *β* in Eq. (1)-(3) and the delay between arterial and venous dilations are governed by the *τ* parameter in Eq. (3) (a longer *τ* correlates to a longer delay).

### 4.7 Investigating BOLD contributions from individual vascular compartments

Our simulations of the BOLD activation profile across cortical depths revealed a mid-depth “bump”, or a local maximum, overlapping with the middle layers in both rVAN1 and rVAN2 (**Fig. 6A**). When simulating by compartment, a similar bump appeared only in the capillary compartment. For reference, we also show the baseline capillary volume showing a similar bump in the middle cortical layers.

Because the reconstructed VAN models lack large pial vessels, they by themselves cannot be used to investigate the effects of draining veins running along the cortical surface on the BOLD cortical-depth profiles. To address this, we grafted a single large vein on the pial surface (see **Fig. 6B**) and tested to what extent it influenced the resulting cortical-depth profile of the BOLD response. As expected, the large pial vein caused a sharp increase in the magnetic field offsets concentrated primarily at the pial surface, as shown in **Fig. 6B**.

We noted our choice of a uniform CMRO_2_ across cortical depth caused a depth-varying baseline oxygenation in the VAN models as visualized in **Fig. 6C**. This choice may, in turn, impact the BOLD profile, contributing to the observed “bump” so we also simulated the layer-BOLD profile after tuning a depth-dependent CMRO_2_ value to achieve a relatively uniform baseline oxygenation profile (**Fig. 6C**, right). We chose a linearly decreasing CMRO_2_ model as the simplest assumption, with CMRO_2_ highest in shallow layers near to the cortical surface and lowest in deep layers near to the white matter, which is also in line with previous estimations from optical measurements (Mächler et al. 2022).

We then tested the cortical-depth profiles of the BOLD response using these two CMRO_2_ models with and without the large pial vein grafted onto the pial surface. As expected, the cortical-depth profiles of the BOLD response amplitude was unaffected beyond the superficial cortical depths and little effect was observed 350 μm below the CSF interface (**Fig. 6D**–**E** for rVAN1 and rVAN2, respectively). When moving from the uniform CMRO_2_ model to the depth-dependent CMRO_2_ model, we observed a more pronounced decay of BOLD amplitude with cortical depth yet the middle-layer “bump” in the BOLD profile was still seen using both CMRO_2_ models (**Fig. 6D**–**E** for rVAN1 and rVAN2, respectively).

Lastly, we observed low oxygenation values in some capillaries and veins but acknowledge this is a direct consequence of the OEF value used, which again was taken from previous measurements (see *Methods*). We note that this OEF value translates to ∼51% reduction in pO_2_ between the arteries (102 mmHg pO_2_) and the veins. While the large variability in capillary oxygenation seen here may seem counter-intuitive, this high degree of heterogeneity and this broad range of values were previously demonstrated in similar VAN modeling studies and shown to be similar to experimental data (Hartung, Badr, Moeini, et al. 2021; Gould et al. 2017).

## 5 Discussion

Here we extended biophysical VAN simulations to the scale of the human cerebral cortical thickness to enable direct comparisons and determine how vascular anatomical differences between rodents and humans impact the BOLD fMRI response to neuronal activity. We introduced several methodological advancements to make a human VAN simulation possible. For *anatomical input data*, we generated a synthetic human VAN with geometric and topological properties derived from human histology data. For *physiological input data*, we extrapolated single-vessel vascular responses to neuronal activity measured from rodents to the thicker human cortex. To overcome the computational burden of larger human VAN sizes, we adapted a scalable oxygenation model. Combined, our framework enabled testing our hypothesis that the differences in vascular architecture between mice and humans would lead to observable differences in their respective BOLD responses. In our analyses, we considered differing geometric features (distances and densities) and topological features (connectivity and branching) in mice and humans.

We found unexpected similarities between species including minimal arterial branching (the MAB principle) observed in both mouse microscopy data (Blinder et al. 2013; Gagnon et al. 2015) and human histology data (Duvernoy et al. 1981). This unexpected asymmetry between branches off diving arterioles and ascending venules—a topological feature—appears necessary for obtaining strong BOLD response amplitudes in synthetic VANs. Using a circuit-theory analogy, these branches act like parallel resistors between the larger vessels (arterioles and venules) and the mesh-like capillary bed. The vascular response to neuronal activity is arterial dilation (lowering resistance), thus baseline arterial resistance must be large to elicit a strong flow response. Having fewer branches off diving arterioles increases baseline arterial resistance and more branches off ascending venules decreases venous resistance that further increases the relative resistance of the arterial compartment. Therefore, the largest baseline arterial resistance amongst these configurations would emerge from fewer branches of diving arterioles and many branches off ascending venules, exactly as observed in the anatomical microscopy data. It may be possible, however, to impose an exaggerated arterial dilation in VANs without MAB to compensate for the lower baseline resistance. As MAB was observed in all anatomical data investigated, and without evidence for larger dilations in regions with more branches off diving arterioles, we conclude that MAB is likely necessary for producing realistic blood flow and BOLD responses in both mice and humans. Aside from the hemodynamic response, these branching properties are likely needed to improve arterial control, ensuring its role in active flow and pressure regulation is maximized.

The implications of MAB for *in-vivo* blood flow regulation are unclear. Because the capillary bed presents the highest resistance to flow within the vascular network (Gould et al. 2017), it is reasonable that there is an asymmetry in the number of arteriolar branches and the number of venular branches that confers higher baseline resistance on the pre-capillary side, and thus more capability to regulate flow through arteriolar dilation. The relatively fewer number of arteriolar branches may also have implications for fine-scale blood flow regulation at the level of individual cortical layers, and the greater number of venular branches may hinder identification of which cortical layer or layers is/are activated based on fMRI contrasts weighted more towards venous signals like BOLD. Insofar as arteriolar branching topology is conducive to blood flow control, our observations are compatible with recent work suggesting that a large portion of blood flow regulation occurs at pre-capillary arterioles at the “arteriole-capillary transition” or ACT zone (Hartmann et al. 2022; Mughal et al. 2023). The fact that mice and humans exhibit this striking feature perhaps hints that there may be similarities in blood flow regulation enacted by pre-capillary arterioles that are conserved between these species. Regardless of whether the MAB property points to commonalities in blood flow regulation across species, the remaining differences in vascular architecture between mice and humans will still likely lead to differences in the timing or shape of the BOLD response between species as seen in **Fig. 2**.

We indeed found species-specific differences in the simulated BOLD responses. For instance, the hemodynamic response to long-duration stimuli differed between species (**Fig. 5**). Although the artery-to-vein ratio is larger in the human VAN—another *topological* feature—resulting in more blood delivered to each vein, contrary to our expectations venous ballooning during long-duration stimulation was smaller and slower in human than in mouse. This may be explained through the greater spacing between arterioles and venules in humans (a *geometric* difference), leading to longer capillary paths and raising the resistance of the capillary compartment. This reduces the relative contribution of arterial resistance to the total resistance of the network (discussed below), causing a reduced pressure redistribution during activation, reducing the passive ballooning in the capillaries and veins (**Fig. 5**). The longer capillary paths also lead to slower pressure redistribution and passive dilation in capillaries and veins in the human sVAN. This reduced amplitude of venous volume response may impact BOLD response dynamics in humans. Our analysis indicates that the human vascular architecture alone will likely result in some hemodynamic differences between species—one can speculate that these architectural differences may result either in compensatory differences in active dilation of arteries, or perhaps in passive dilation of capillaries and veins. We do note, however, that these predictions only account for our current understanding of passive ballooning mechanics and structural properties of the human microvasculature. These species-specific differences between venous ballooning still require validation with *in-vivo* measurements such as single-vessel fMRI in human (Varadarajan et al. 2023; Hartung et al. 2022) and mouse (Yu et al. 2014; 2016).

There are many possible explanations for this reduced venous ballooning in our simulation, and our results do not prove a reduction in ballooning in humans compared to mice. Yet they do suggest that prior intuition about the impact of vascular architecture on hemodynamic responses may not always be correct. Given the many interdependent properties of the model, further simulations are needed to understand the full extent of how vascular architecture influences the BOLD response. Advanced fMRI techniques that report hemodynamics from different vascular compartments may assist determining whether venous dilation during long-duration stimuli differs between humans and mice (Chen and Pike 2009).

The timing of simulated BOLD responses also differed between mice and humans, which may be attributed to the increased length of diving arterioles spanning the cortical thickness. This geometric difference increases the distances of upstream dilation signaling during activation. We tested this theory by extrapolating recorded arteriolar dilations from mouse to the thicker human cortex. Scheme 1, simply assigning dilations from relative cortical depth, achieved faster BOLD timing as mouse rVANs. Scheme 2 accounted for longer upstream signaling distances and resulted in a BOLD response similar to that from the mouse VANs. Schemes 3 and 4 used more elaborate geometrical scaling that accounts for other delays. These Schemes resulted in a slower BOLD response consistent with experimentally measured BOLD fMRI data (Lambers et al. 2020). This BOLD timing difference via dilation Schemes implies increased VAN size and/or vessel path-lengths (geometric effects) may help explain slower hemodynamic responses in humans. This also implies blood rheological properties varying between species (red blood cell size, cardiac frequency, etc.) and vascular topological differences may be insufficient to account for hemodynamic timing differences observed in empirical fMRI data. This is a testable hypothesis where arterial dilation measurements in humans can confirm whether the dilation time-courses indeed match our estimated dilations generated from Schemes 3 and 4.

Our mouse simulations use dilation traces as inputs, measured from anesthetized mice (isoflurane) following electrical forepaw stimulation and generate as outputs BOLD responses, which we in turn compared to the corresponding BOLD responses measured in anesthetized mice. The anesthetic used for both sets of *in-vivo* measurements is known to cause a dose-dependent increase in CBF response amplitude and speed (Masamoto et al. 2009). In addition, a previous study demonstrated an interaction between the level of this anesthetic and the pulse-train interval used for forepaw electrical stimulation and the resulting CBF response (Masamoto et al. 2009), which showed that, for the shortest pulse-intervals tested (50 ms), the CBF response amplitudes and speeds were consistent across anesthetic doses. Fortunately, the pulse-train interval used to generate the *in-vivo* measurements used for our simulations was shorter than this stable region (six pulses with a ∼25 ms interval (Tian et al. 2010)). Thus, we expect that the measured data we used as inputs to and validations of our simulations may also be less influenced by the specific anesthetic dose used.

In addition to considerations related to the depth of anesthesia used for the *in vivo* microscopy and BOLD fMRI measurements in mice, there are also considerations for relating data collected in anesthetized mice to data collected in awake/unanesthetized humans, which somewhat complicates the comparison of interest between mouse and human. Anesthesia affects baseline blood flow (Gao et al. 2017), however different anesthetics have different effects (Franceschini et al. 2010), and the hemodynamic response to neuronal activity is generally believed to be slower in anesthetized animals (Le et al. 2024; Masamoto et al. 2009). These effects will impact the BOLD response timing and amplitude, although we would expect that, if similar arterial dilation traces recorded across the vascular hierarchy were available from the awake mouse, the hemodynamic response—including the BOLD fMRI response—would be faster, therefore we would expect even greater differences between mice and humans, which would further strengthen our claim that mouse and human hemodynamics differ. While our study may not provide any specific insights into these well-known effects of anesthesia, future VAN modeling work may be able to investigate how altered arterial dilations manifest in altered BOLD responses under anesthesia relative to the awake state. Nevertheless, correcting for the effects of anesthesia on the CBF and BOLD responses may be feasible, and the largest change in the responses expected in an awake preparation would be a smaller and faster CBF (and thus BOLD) response in the mouse VAN, which would potentially translate into a somewhat smaller and faster BOLD response across all Schema tested.

While our BOLD responses from human sVANs are arguably realistic, given that the data used for the inputs to our modeling framework are unavailable in humans our simulated responses result from educated guesses that rely on several assumptions. We assume the arterial signaling mechanisms in mice and humans lead to equivalent upstream dilation propagation velocity through the arterial tree. Without evidence to the contrary, we consider this a safe assumption. We also assume dilations originate in the bottommost cortical layer, Layer VI, although the dilation recordings we use do not differentiate individual layers. In reality, neuronal activity and associated arterial dilations may originate in Layer IV and spread upwards and downwards simultaneously (Fracasso et al. 2016). We chose not to implement this to avoid data overfitting. Another key assumption is that the dilation does not stop after stimulation ends. Instead, in our simulations the full dilation sequence is carried out, causing some vessels to peak at ∼5 s, long after stimulation ends, again to avoid overspecification and overfitting. Recent progress towards non-invasive “single-vessel fMRI” could revisit these assumptions with the potential to measure response timing in humans (X. Chen et al. 2021; Varadarajan et al. 2023).

Additionally, our VAN simulations are isolated from oxygen supply/drainage and from outside or below the VAN field-of-view. Our no-flux boundary condition, however, maintains moderate oxygenation near these interfaces. This assumption is similar to assumptions made in prior studies (Gould et al. 2017; Schmid, Tsai, et al. 2017; Linninger et al. 2013; Hartung, Badr, Moeini, et al. 2021; Genois et al. 2021; Reichold et al. 2009; Damseh et al. 2021; Báez-Yáñez and Petridou 2024; Báez-Yáñez et al. 2025; Lorthois et al. 2011), however in future work the these boundary conditions could be validated by synthesizing or reconstructing larger VANs and activating only the central portion (Pfannmoeller et al. 2021). Moreover, we chose a linearized oxyhemoglobin dissociation model to decrease the computational burden, although efficient nonlinear extensions using fast Fourier-based solvers could be applied in future work (Ventimiglia and Linninger 2023; Linninger et al. 2024).

We implemented a simple compliance model based on the Balloon Model. While simpler than proposed viscoelastic models (Park et al. 2020; Pfannmoeller et al. 2021; Krieger et al. 2012), our approach captures the same biophysical relationships. The parameter *β* modulates the dilation amplitude during pressure changes as the elasticity term in viscoelastic models (Pfannmoeller et al. 2021). The parameter *τ* controls the dilation speed as a viscosity parameter. In future work more advanced models may provide insight into BOLD nonlinearity and how dynamics change across stimulus configurations (Pfannmoeller et al. 2021).

We impose active dilations only on arterioles and passive dilations on capillaries and veins. However, active control mechanisms, such as pericyte control, may regulate blood flow in capillaries, although pericyte-mediated active dilation at the time scales of functional hyperemia remains controversial (Berthiaume et al. 2018; Cai et al. 2018; Hartmann et al. 2022). Inclusion of pericyte-driven capillary dilation would be straightforward in our framework if the spatial distribution and temporal coordination were known.

Our oxygen model does not change the metabolic rate constant with local activity. Our first-order model does, however, modulate the metabolic rate according to local oxygen tension: as the supply increases during activation, metabolism follows (see *Supplementary Notes*). Nevertheless, we do not expect that a different model would substantially affect our observations or conclusions. Moreover, we calibrated both the simulations based on human and mouse VANs to the same baseline perfusion rate (100 mL/100g/min) to isolate vascular architecture as the main difference between species; these simulations can readily be performed with a more realistic baseline perfusion for human (∼60 mL/100g/min (Juttukonda et al. 2021; Liu and Brown 2007; Buxton 2009)). This higher CBF led to the use of a higher CMRO_2_ to achieve the expected 30% OEF in our human model. Nevertheless, this change in baseline perfusion, one of the only two calibrations used for our hemodynamic simulations (see *Methods*), will not affect our simulation results with the potential exception of the amplitude of the initial dip (discussed next). However, a higher baseline CBF will be accompanied by a concomitant reduction in baseline CMRO_2_. This is because our calibration adjusts each VAN simulation to achieve the target 30% OEF, as detailed in the *Methods* section. Thus, any reduction in baseline CBF and thus CMRO_2_ would generate the same baseline values for oxygen tension in the vasculature and tissue, and would in turn generate equivalent oxygenation patterns (in space and time), resulting in unchanged BOLD responses.

We observed a higher oxygen content in pial veins than diving veins deeper in the cortex (see **Fig. 6B-C**). This effect has also been observed empirically with optical imaging (Li et al. 2019). This may be counter-intuitive under the assumption that oxygen content decreases steadily in blood traveling from the arteries to the veins. Here, however, the oxygen content in ascending and pial veins is a function of the vascular connectivity patterns. The veins in superficial cortical depths are more oxygen-rich due to the higher arterial density and higher oxygen in the capillaries at superficial depths. This results in lower oxygen content in deeper parenchymal veins that combines with blood from oxygen-rich capillaries while traveling to the surface veins, increasing the oxygen content of the more superficial veins. This agrees with data suggesting oxygen content in veins increases with diameter (Gagnon et al. 2015; Sakadžić et al. 2014) and with decreasing branch order (Sakadžić et al. 2014; 2010).

We also observed an initial dip in the simulated BOLD response in rVANs that was nearly absent in sVANs (**Fig. 3**). The “elusiveness” and prevalence of the initial dip in BOLD fMRI data is still debated in the fMRI community, so a full investigation is outside the scope of this work. We attributed this initial dip to slow blood velocity in the arterioles in the rVANs due to the many disconnected pial arterial inlets and venous outlets. We verified this hypothesis in a post-hoc analysis consisting of (*i*) connecting all terminals to a single inlet and outlet, increasing the velocity in many arterioles, which eliminated the initial dip in subsequent simulations, and (*ii*) reducing the velocity in arterioles, venules, and capillaries, which generated an initial dip (See *Supplementary Notes*). Also, the lower velocity in arterioles in sVANs with more branches off diving arterioles (**Fig. 4E**) generated an initial dip. Intuitively, while vessels are expanding during initial dilation, blood outflow from expanding vessels will transiently decrease to fill the volume—i.e., vessel expansion leads to a “transverse” flow component that causes a reduction in outflow due to mass conservation (Mandeville et al. 1999). This leads to longer residency time, increased oxygen extraction, and a dip in blood oxygenation in downstream vessels.

Vascular architecture is known to vary across cortical regions (Tsai et al. 2009). While a key potential application of the VAN framework would be to investigate how hemodynamics vary across brain regions, the recordings of arterial dilations that serve as inputs to our simulations are currently only available for the somatosensory cortex, limiting the application to other regions. Moreover, either accurate reconstructions of the full vascular architecture, or the statistical distributions of geometry and topology needed for vascular synthesis, are not currently available for other brain regions. However, if these anatomical and physiological data, as well as the corresponding BOLD responses needed for validation, were collected in other regions these data can be readily fed into our framework to further investigate whether the relationship between microscopic vascular dynamics and the measured BOLD fMRI responses is conserved across the brain. This will be challenging, partly because it is well-known that BOLD response timing varies within cortical areas, including primary sensory cortex in humans (Polimeni and Lewis 2021; Gomez et al. 2024), and so we do not expect that the timing of our simulated BOLD response simulated at one cortical location should match that of measured BOLD responses across the entire cortical area. Also, BOLD response timing varies with the specifics of the stimulus or task configuration including duration (as mentioned above) and intensity (J. E. Chen et al. 2021), therefore microvascular and BOLD responses should always be measured with the same stimulus/task. However, if both the required anatomical data—either full reconstructions or statistical distributions of the vascular geometry and topology—and the required physiological data—including single-vessel arterial dilation responses and the corresponding BOLD fMRI responses to neuronal activity—were available from other brain regions, the framework presented here could be used to extend VAN-based modeling beyond primary somatosensory cortex. Furthermore, our human VAN used geometric parameters from the collateral sulcus of the temporal lobe (Cassot et al. 2006), a region not commonly studied in BOLD fMRI experiments. However, the ink-injection for vascular staining performed well in this region, making it a suitable candidate for statistical analysis and reconstruction (Cassot et al. 2006). If the statistics were made available from other human brain regions, an sVAN could readily be generated with these statistics. Additionally, these statistics could confirm the MAB principle as a general property across brain regions.

More broadly, our framework would benefit from more microvascular data in humans, which are more challenging to obtain compared to mice for practical reasons. Moreover, although the available human microvascular anatomical data are highly valuable, they were acquired and analyzed using different procedures than those used for the mouse data, leading to differing anatomical biases. For example, in **Fig. 1A** the vessel diameter histograms suggests that human capillary diameters are ∼2–3 μm larger than those in mice even though human red blood cells (RBCs) are only ∼1 μm larger. Furthermore, the vessel segment length distributions are unexpectedly similar between humans and mice, which would suggest that the two species have comparable vascular densities, assuming similar branching patterns. If microvascular data in humans and mice could be obtained in homologous brain areas with similar acquisition and analysis techniques, we would be able to compare with greater certainty the anatomical differences.

Our investigation of compartment-specific BOLD cortical-depth profiles indicated that the main contributor is the capillary compartment—the BOLD response profile closely reflected capillary blood volume. The arterial and venous compartments mainly affect the profile in locations near the pial surface. This was further confirmed in our BOLD simulations incorporating a large vein grafted on the pial surface, which we interpret being caused by the magnetic field generated by these large pial vessels that falls off inversely with distance, limiting the reach of the magnetic field perturbations generated by these vessels.

The implications of vascular differences between mouse and human are a source of future investigations. For example, humans have 4× fewer diving arterioles, 23.4× fewer ascending venules, larger vessel diameters, and reduced capillary density than mice. This may minimize overall blood volume while delivering sufficient nutrients, as previously suggested (Murray 1926). This relative sparsity of large vessels may also introduce disadvantages; for instance, the larger capillary domain fed by a single diving arteriole potentially reduces the specificity of blood flow regulation accompanying neuronal activation. Conversely, the nearly flipped artery-to-vein ratio in humans increases the relative arterial density, partly restoring control. These two features may balance each other to create efficient blood delivery to the much larger human cortex while achieving sufficient spatial and temporal specificity of the hemodynamic response.

Additionally, the thicker human cortex and the longer distance traveled by the upstream propagation of dilation during activation may lead to more localized responses in humans compared to mice. For example, if upstream dilation stops propagating after stimulus offset, a short-duration (e.g. 2-s) stimulus may only dilate the deepest arterioles in humans yet dilate arterioles across the entire cortical thickness in mouse. Identifying such differences in hemodynamic control in humans would help with interpretation of human fMRI data.

Despite species differences, mice are the most common experimental model for human neurovascular coupling due to the many similarities in their cerebrovascular architecture and neuronal responses to sensory stimulation. This includes shared microvascular hierarchy (pial arteries, diving arterioles, capillaries, ascending venules, pial veins) and the relationship between vascular and functional architectures. Yet mice are available for invasive experimentation not suitable for humans, and insights from mice appear to explain many aspects of fMRI data in humans (Gagnon et al. 2015). It is still important, however, to be mindful of observations in mouse that may not generalize to humans. More high-resolution anatomical and functional data are needed in humans to validate our findings, provide more accurate inputs, and confirm in which ways the mouse experimental model can be used to explain human fMRI.

As noted above, many studies over the past two decades have established that blood flow regulation in the brain is controlled over much finer spatial and temporal scales than what was expected at the advent of fMRI (Devor et al. 2003; Nizar et al. 2013; Chen et al. 2011; Cho et al. 2022; Poplawsky et al. 2015; Boido et al. 2019). Unfortunately, the microvascular responses providing the most veridical representation of neuronal activity occur at spatial and temporal scales finer than the imaging resolutions that can be achieved even with modern fMRI technologies (Polimeni and Wald 2018; Hartmann et al. 2022; Drew 2019; Fukuda et al. 2021). Therefore, human fMRI may still have untapped potential and may be capable of more accurate measurement of the neuronal activity of interest. Uncovering this information would require models that relate the observable BOLD fMRI signals to unobservable microscopic hemodynamic changes. Unfortunately, fully inverting this relationship to infer neuronal activity from the BOLD response, or “deconvolving” the hemodynamic response from the fMRI data, remains challenging. A more complete understanding of the vascular filter, however, would likely improve the inference of neuronal activity from fMRI data. Future applications of the human VAN modeling framework may therefore be able to improve the localization of neuronal activity at spatial and temporal scales below the imaging resolution of current human fMRI.

By synthesizing and simulating the first human VAN, we were able to compare how vascular anatomical differences between mice and humans impact the BOLD response. The seemingly most impactful difference is the larger size of the human VAN. Furthermore, the artery/vein branching asymmetry in both species implies that the topological similarities between mouse and human outweigh the differences. Our findings suggest that it is important to consider the differences in vascular architecture between mouse and human brains when interpreting human BOLD fMRI data. As more advanced fMRI contrasts are developed, the VAN modeling framework can be extended to these as well, to provide new insights into how microvascular dynamics influence human fMRI data, again bridging between fMRI and ground-truth measures from optical microscopy, enabling a more accurate interpretation of human fMRI data.

## 6 Data and Code Availability

Source code and data that support the findings of this study will be posted to an open-source repository (github.com) upon acceptance of this manuscript.

## 7 Author Contributions

G.A.H.: Conceptualization, Methodology, Software, Formal Analysis, Investigation, Data Curation, Visualization, Writing—Original Draft & Review & Editing, and Funding Acquisition; A.J.L.B.: Methodology, Writing—Review & Editing; S.S.: Conceptualization and Writing—Review & Editing; A.L.: Writing—Original Draft & Review & Editing; D.A.B.: Conceptualization, Writing—Review & Editing; J.R.P.: Conceptualization, Methodology, Writing—Review & Editing, Supervision, Resources, and Funding Acquisition.

## 9 Acknowledgements

We gratefully acknowledge funding from the NIH (NIBIB grants P41-EB030006, R01-EB019437, and R01-EB032746), from the *BRAIN Initiative* (NIH NIMH grants R01-MH111419 and F32-MH125599, NIH NINDS grant U19-NS123717), from the CIHR (grant MFE-164755), and from the Athinoula A. Martinos Center for Biomedical Imaging. We thank Dr. Joerg Pfannmoeller for his assistance and input during the initial development of the simulation framework. We thank our colleagues Drs. David Kleinfeld, Xiaojun Cheng, Divya Varadarajan, Jingyuan Chen, Jean Chen, Louis Gagnon and Anna Devor for their helpful feedback.

